# Molecular mapping of transmembrane mechanotransduction through the β1 integrin-CD98hc-TRPV4 axis

**DOI:** 10.1101/617092

**Authors:** Ratnakar Potla, Mariko Hirano-Kobayashi, Hao Wu, Hong Chen, Akiko Mammoto, Benjamin D. Matthews, Donald E. Ingber

## Abstract

One of the most rapid (< 4 msec) transmembrane cellular mechanotransduction events involves activation of Transient Receptor Potential Vanilloid 4 (TRPV4) ion channels by mechanical forces transmitted across β1 integrin receptors in endothelial cells, and the transmembrane Solute Carrier family 3 member 2 (CD98hc) protein has been implicated in this response. Here, we show that β1 integrin, CD98hc, and TRPV4 all tightly associate and co-localize in focal adhesions where mechanochemical conversion takes place. CD98hc knock down inhibits TRPV4-mediated calcium influx induced by mechanical forces, but not by chemical activators, thus confirming the mechanospecificity of this signaling response. Molecular analysis revealed that forces applied to β1 integrin must be transmitted from its cytoplasmic C-terminus to the CD98hc cytoplasmic tail, and from there to the ankyrin repeat domain of TRPV4 in order to produce ultra-rapid, force-induced, channel activation within the focal adhesion.

Graphical abstract

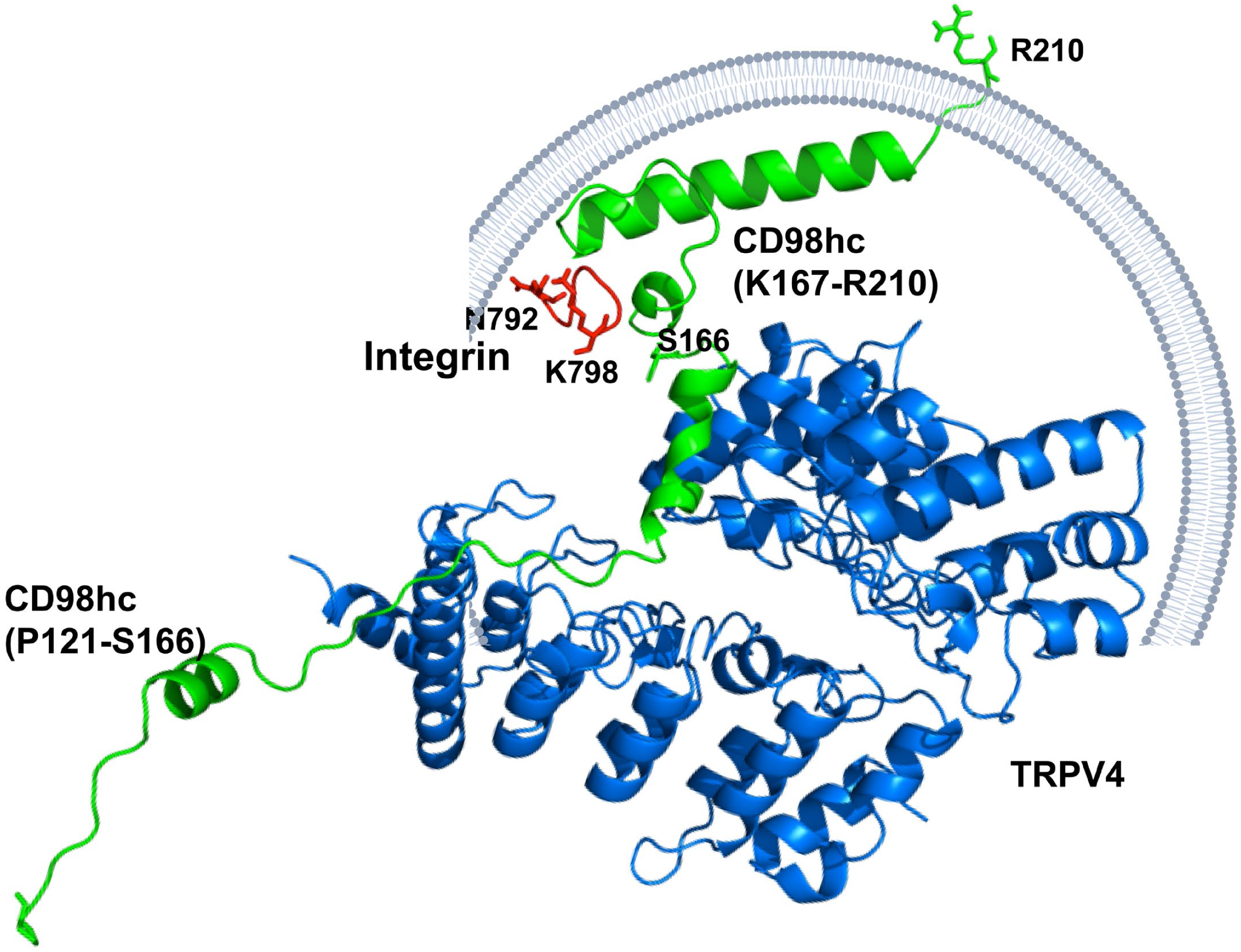

**SUMMARY:** A direct path of mechanical signal transfer between β1 integrin, CD98hc, and TRPV4 channels is identified that mediates ultra-rapid transmembrane mechanochemical conversion within focal adhesions.

## INTRODUCTION

Cellular mechanotransduction is critical for maintenance of cellular growth homeostasis as well as tissue development, and deregulation of this process results in pathogenesis of numerous human diseases (Ingber, 2003). Cell surface integrin receptors sense forces transmitted across the extracellular matrix (ECM)(Wang et al., 1993) and one way they induce mechanochemical transduction is by transmitting these stresses across the cell surface and inducing deformations in proteins, such as talin and vinculin which form the cytoskeletal backbone of the focal adhesion (Choquet et al., 1997). An alternative transmembrane mechanochemical conversion mechanism involves mechanosensitive TRPV4 ion channels on the surface membrane of endothelial cells, which are activated within 4 milliseconds after mechanical forces are transmitted from the ECM to cell surface β1 integrin receptors (Matthews et al., 2010). This response is physiologically and clinically relevant as TRPV4 channels mediate cyclic strain-induced endothelial cell reorientation (Thodeti et al., 2009) and TRPV4 activation restores normal angiogenesis in tumors by modulating the Rho/Rho kinase pathway (Adapala et al., 2016). Activation of TRPV4 by physical breathing motions also contributes to development of pulmonary edema, and drugs that block TRPV4 channel activity prevent pulmonary vascular leakage in a human Lung Alveolus Chip *in vitro* (Huh et al., 2012), as well as development of cardiogenic pulmonary edema in vivo (Thorneloe et al., 2012).

While application of mechanical force to β1 integrins results in almost immediate activation of calcium influx through TRPV4 channels in endothelial cells, the specific path of force transfer from β1-integrin to TRPV4 remains unknown. Past use of single chain integrin truncation mutants revealed that the terminal 6 amino acid residues at the carboxy terminus of the β1 integrin cytoplasmic domain are required for almost immediate induction of calcium influx through TRPV4 when forces are applied directly to the extracellular domain of an engineered, single chain, β1 integrin on the surface of endothelial cells using magnetic tweezers (Matthews et al., 2010). This portion of the β1 integrin C-terminus is also required for adhesion strengthening and cytoskeletal tension-dependent fibronectin fibrillogenesis (Féral et al., 2007) that are mediated by binding of integrin to the single pass transmembrane CD98hc glycoprotein, which also associates with various other multi-pass transmembrane amino acid transporters through its extracellular domain (Kolesnikova et al., 2001; Zent et al., 2000). In a past magnetic tweezer study, CD98hc localized to focal adhesions formed at the site of force application to integrins and siRNA knock down of CD98hc reduced the adhesion strength of the bound β1 integrins on the endothelial cell surface (Matthews et al., 2010). While these studies implicate a role for CD98hc in β1 integrin-dependent stimulation of local calcium influx through TRPV4 ion channels, it remains unclear whether it is directly involved in transferring mechanical forces between integrin and TRPV4 or if it acts indirectly.

TRPV4 is a multi-pass transmembrane protein that forms as a homo- or hetero-tetramer with other TRPV family proteins (White et al., 2016). The TRPV4 protein consists of a cytoplasmic amino terminus, six transmembrane domains, and a cytoplasmic carboxy terminus. The cytoplasmic amino terminus of TRPV4, which contains a N-terminal tail, proline rich domain (PRD), and six ankyrin repeat (AR) domains, has been shown to bind to cytoskeletal proteins, such as actin and tubulin (Goswami et al., 2010). TRPV4 transmembrane domains also interact with cholesterol and play a role in its localization to lipid rafts (Kumari et al., 2015), whereas the cytoplasmic carboxy terminus plays a role in regulation of TRPV4 folding, maturation and trafficking (White et al., 2016).

In this study, we set out to determine if the cytoplasmic tail of β1 integrin, CD98hc, and TRPV4 bind directly to each other, and if so, to identify the specific domains and residues in these molecules that mediate transfer of mechanical forces from β1 integrin through CD98hc to TRPV4 that are responsible for its activation. Here, we report the discovery of specific amino acid (aa) residues located within the 50 aa (K160 – C210) high homology (HH) domain of CD98hc that bind the NPKY motif found within the β1 integrin cytoplasmic tail, as well as other residues within CD98hc’s cytoplasmic tail that bind to the AR domain of TRPV4. Importantly, we also demonstrate that while CD98hc is required for activation of TRPV4 by mechanical stimulation, it is not necessary for its activation by chemical inducers. This new understanding of the molecular path of direct mechanochemical conversion at the plasma membrane involving β1 integrin, CD98hc and TRPV4 may aid in the future development of more specific and effective mechanotherapeutics that target diseases, such as pulmonary edema, where mechanotransduction through TRPV4 plays an important role.

## RESULTS

### β1 integrin, CD98hc, and TRPV4 tightly associate in focal adhesions

Deletion of the last 6 aa within the cytoplasmic tail of β1 integrin inhibits mechanical force-induced calcium signaling in bovine capillary endothelial cells and human dermal microvascular endothelial cells (Matthews et al., 2010). When human embryonic kidney (HEK) 293T cells were transfected with wild type β1 integrin (β1) or single chain β1 integrin mutant constructs that lacked either 6 aa in the juxtamembrane region (β1Δ1) or the last 6 aa at the cytoplasmic tail β1Δ5 (Supplementary Figure S1A), only the β1 and the β1Δ1 mutant containing the intact cytoplasmic terminus co-immunoprecipitated with both CD98hc and TRPV4 (Figure 1A). Moreover, when similar studies were carried out in human umbilical vein endothelial (HUVE) cells using anti-TRPV4 antibody, endogenous β1 integrin and CD98hc co-precipitated (Figure 1B). Similarly, antibodies directed against either β1 integrin or CD98hc resulted in co-precipitation of complexes containing TRPV4 (Figure 1C).

**Figure 1.**
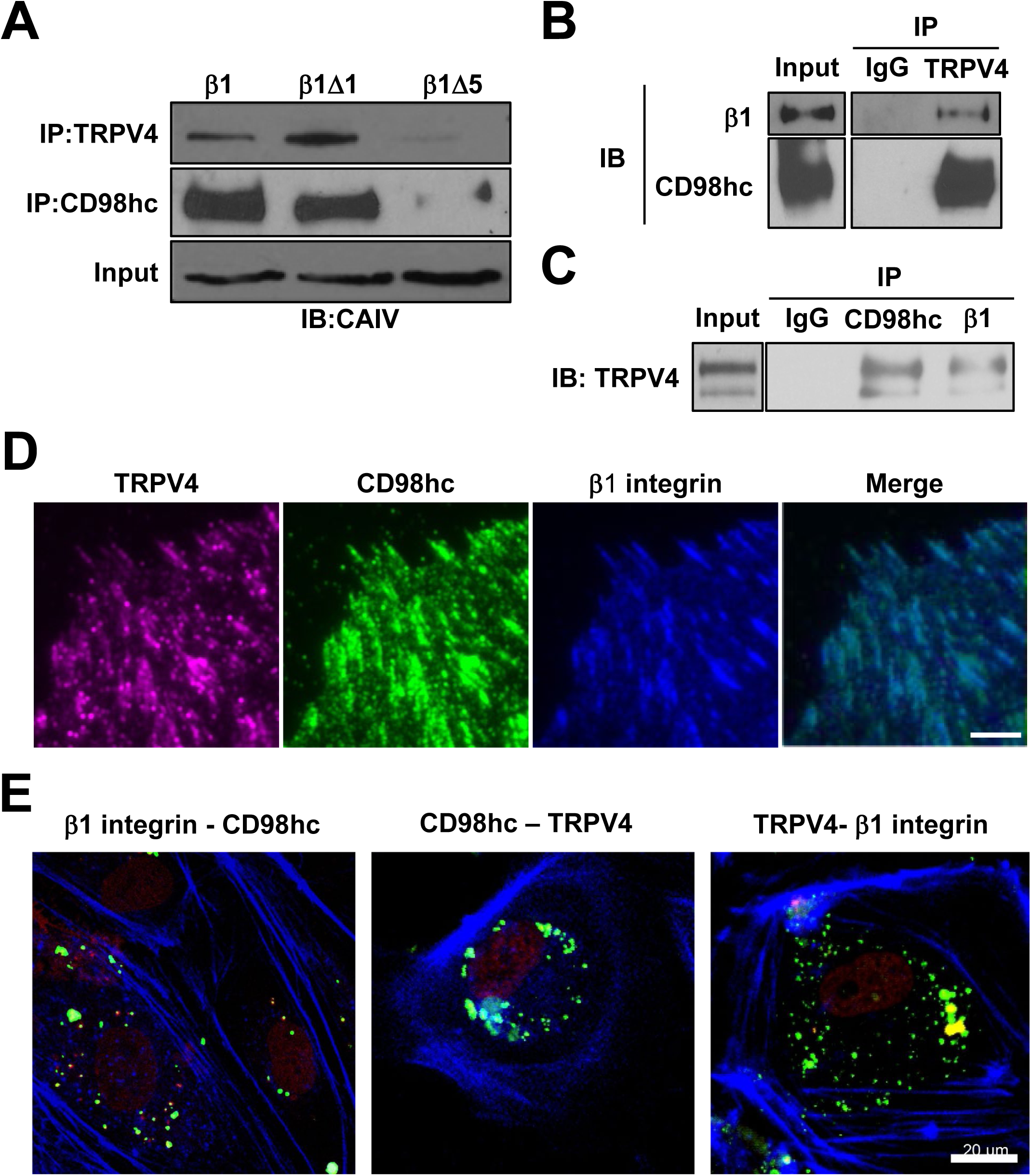
β1-integrin, CD98hc, and TRPV4 co-associate in focal adhesions in adherent cells. (**A**) Immunoblots showing β1-integrin (full-length, β1Δ1, β1Δ5) immunoprecipitated with TRPV4 and CD98hc antibodies from total cell lysate of HEK293T cells transfected with CA-LDL−β1 (β1), CA-LDL−β1Δ1 (β1Δ1), and CA-LDL−β1Δ5 (β1Δ5) chimerae. Immunoblots showing β1-integrin and CD98hc immunoprecipitated with TRPV4 antibody or IgG from total cell lysate of HUVE cells (**B**) and TRPV4 immunoprecipitated with CD98hc and β1-integrin antibodies from total cell lysate of HUVE cells (**C**), with IgG used as a control. (**D**) TIRF micrographs of control HUVE cells showing localization of TRPV4 (magenta), CD98hc (green), β1-integrin (blue) in focal adhesions, as well as a merged image of all signals (right) (bar, 5 μm). (**E**) Confocal micrographs of HUVE cells showing proximity ligation foci (green) between endogenous CD98hc and endogenous TRPV4 (CD98hc-TRPV4), overexpressed CD98hc and endogenous TRPV4 (FLAG-TRPV4), overexpressed CD98hc and endogenous β1-integrin (FLAG-β1) (bar, 20 μm)

Analysis of protein distribution using total internal reflection fluorescence (TIRF) microscopy confirmed that endogenous TRPV4 colocalizes with both CD98hc and β1 integrin within spontaneously formed focal adhesions along the basal membrane of adherent HUVE cells (Figure 1D). When a Proximity Ligation Assay was carried out that reliably detects specific protein-protein interactions within a 40 nm radius of the target protein (Fredriksson et al., 2002), we found that while CD98hc and TRPV4 proteins associated closely with each other along the entire basal and apical cell membranes, tight interactions between CD98hc and β1 integrin were limited to the basal cell surface (Figure 1E) where focal adhesions are located (Figure 1D). This was accomplished by overexpressing a FLAG-tagged peptide containing 1-210aa of CD98hc (Supplementary Figure S1B) in HUVE cells and performing the proximity ligation assay using anti-TRPV4 or anti-β1 integrin antibody in combination with anti–FLAG antibodies. Taken together, these findings suggest that β1 integrin, CD98hc and TRPV4 tightly co-associate with each other in common basal focal adhesions where they co-localize within human endothelial cells.

### CD98hc physically couples β1 integrin to TRPV4 in focal adhesions

To determine whether CD98hc mechanically links β1 integrin to TRPV4, we knocked down CD98hc in HUVE cells using siRNA (Supplementary Figure S2A). This greatly reduced the amount of both CD98hc and TRPV4 within focal adhesions, although TRPV4 staining could still be detected at other regions of the cell membrane (Figure 2A). This effect was specific as knocking down CD98hc expression had no effect on recruitment of either integrin or vinculin to the focal adhesion (Supplementary Figure S2B). It also did not interfere with the structural function of focal adhesions, as indicated by the absence of any suppression of cell migration in scratch wound assays (Supplementary Figure S2C). Therefore, CD98hc is not required for focal adhesion formation, integrin recruitment to these sites, or cell movement. On the other hand, knockdown of CD98hc reduced the ability of β1 integrin and TRPV4 to co-associate with each other by almost 80%, as determined using a co-immunoprecipitation assay (Figure 2B, C). These results indicate that CD98hc physically couples β1 integrin to the subset of TRPV4 molecules that are located within the focal adhesions.

**Figure 2.**
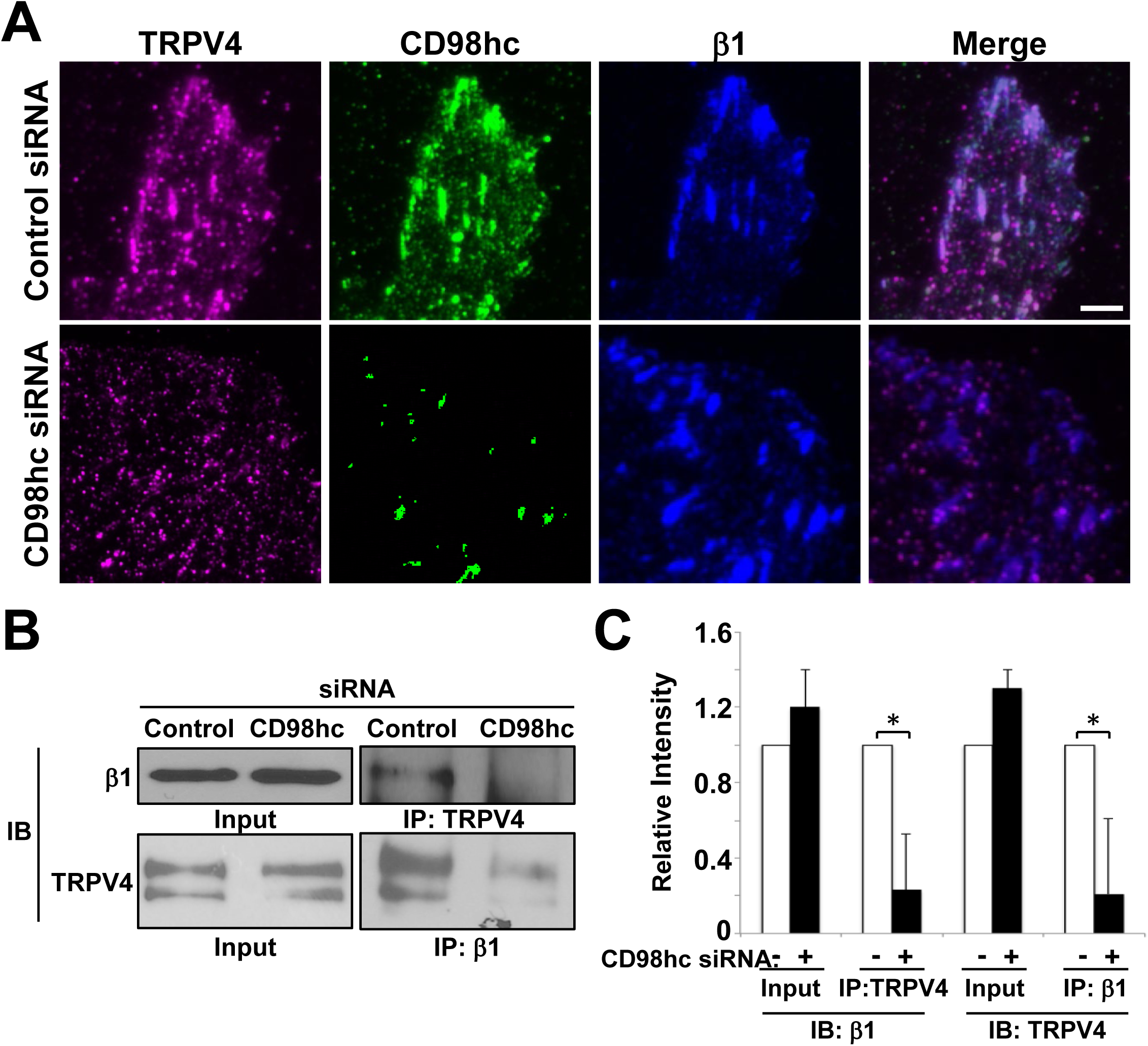
CD98hc is not required for focal adhesion formation but is needed for recruitment of TRPV4. (**A**) TIRF micrographs of control HUVE cells or HUVE cells treated with CD98hc siRNA showing localization of TRPV4 (magenta), CD98hc (green), and β1-integrin (blue) in focal adhesions, as well as a merged image of all signals (right) (bar, 5 μm). (**B**) Immunoblots showing β1-integrin immunoprecipitated with TRPV4 antibody or TRPV4 immunoprecipitated with β1-integrin antibody from lysates of HUVE cells treated with control or CD98hc siRNA. (**C**) Relative intensities of signals from the blots in panel B (all data shown are mean +/- SEM., * p<0.05).

### CD98hc HH domain mediates binding between β1 integrin and TRPV4

We then set out to map the molecular path by which CD98hc mediates force transfer from β1 integrin to TRPV4. The highly conserved HH domain of CD98hc consists of 50 aa (160-210 aa) spanning its transmembrane, extracellular juxtamembrane, and intracellular juxtamembrane regions (Supplementary Figure S3A). When HEK293T cells were transfected with intact single chain β1 integrin (Supplementary Figure S1A) and either full length CD98hc or CD98hc lacking the HH domain [CD98hc(ΔHH)] (Figure 3B, Supplementary Figure S3A), β1 integrin and TRPV4 only co-immunoprecipitated with the intact molecule (Figure 3B). Thus, the HH domain of CD98hc appears to mediate its binding to β1 integrin, and it is required for co-association with TRPV4; however, it remains unclear whether TRPV4 associates with this same domain or if it interacts with a different region of the CD98hc molecule.

**Figure 3.**
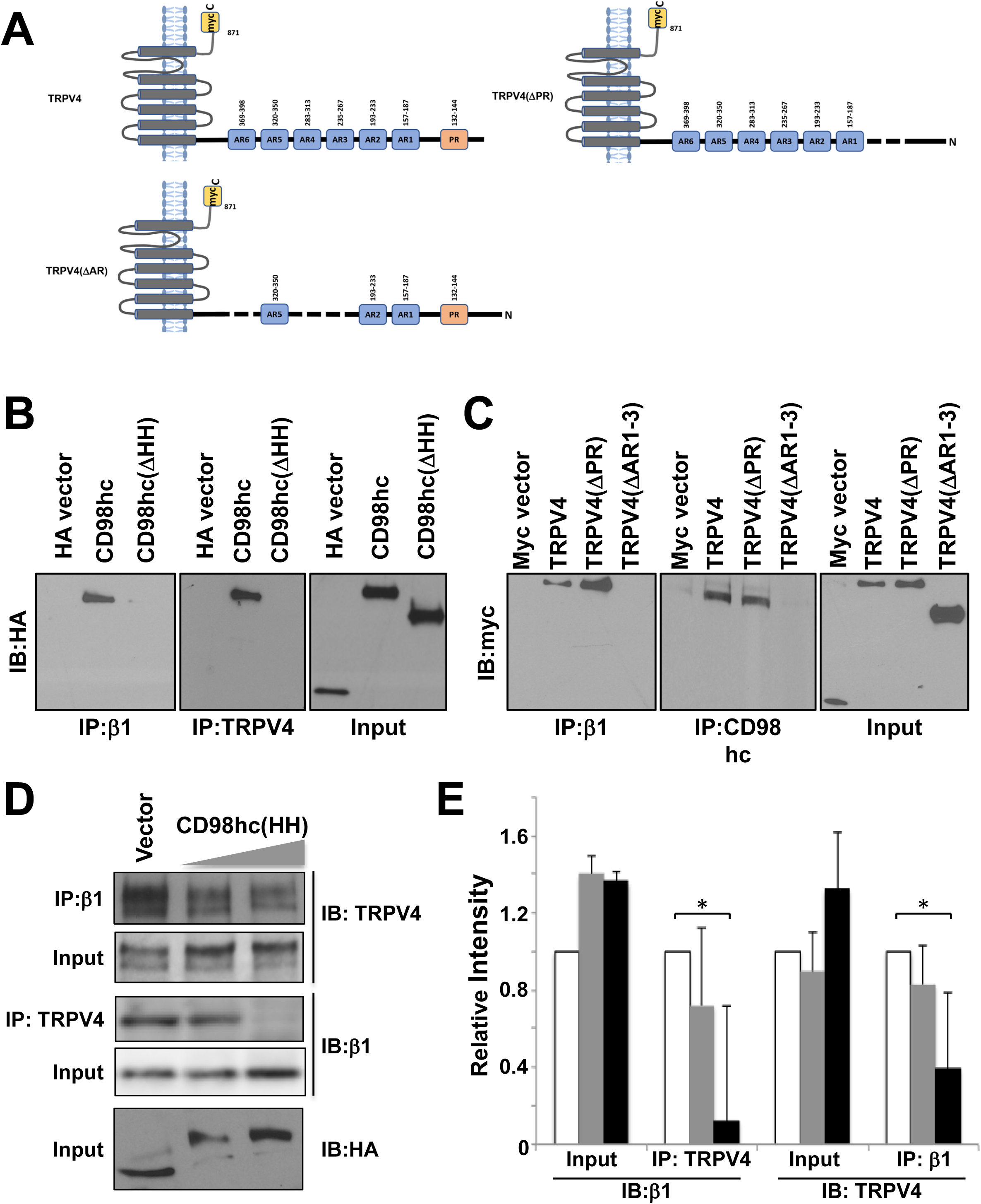
CD98hc HH domain mediates the binding between β1-integrin and TRPV4. (**A**) Schematic diagram of TRPV4 constructs showing transmembrane, proline rich (PR), and ankyrin repeat (AR, blue) domains, as well as myc (yellow) tags. Immunoblots showing HA-tagged CD98hc constructs immunoprecipitated with β1-integrin (left) or TRPV4 (middle) antibody in lysates from HEK293T cells transfected with indicated DNA constructs (**B**), myc-tagged TRPV4 constructs immunoprecipitated with β1-integrin (left) or CD98hc (middle) antibody in lysates from HEK293T cells transfected with indicated DNA constructs (**C**), and β1-integrin and TRPV4 immunoprecipitated with β1-integrin and TRPV4 antibodies, respectively, in lysates from HEK293T cells transfected with increasing amounts (2.5, 5 µg) of the CD98hc(HH) domain construct (**D**). (**E**) Histogram showing the densitometric quantification of immunoblot signals from experiments described in **D**, where bars are vector control (white), CD98hc(HH) low (grey), and CD98hc(HH) high (black).

To better understand this mechanism, we next attempted to identify regions within TRPV4 that mediate its association with CD98hc by genetically engineering TRPV4 mutants in which the transmembrane [TRPV4(N)] or C-terminal regions of the molecule [TRPV4(ΔC)] are deleted (Supplementary Figure S3C). Removing these domains did not alter TRPV4’s ability to coimmunoprecipitate with either β1 integrin or CD98hc (Supplementary Figure S3D), and deletion of the proline-rich domain in the TRPV4 N-terminus [TRPV4(ΔPR)] also had no effect (Figure 3C). However, deletion of three AR domains (235-267aa, 283-313aa, and 369-398aa) in the N-terminus of TRPV4 [TRPV4(ΔAR)] (Figure 3A) was sufficient to completely inhibit association of TRPV4 with both CD98hc and β1 integrin (Figure 3C). Furthermore, when increasing amounts of the CD98c HH domain [CD98hc (HH)] were overexpressed (Supplementary Figure S3A) in HEK293T cells, it acted as a dominant negative protein and decreased the amount of β1 integrin that co-immunoprecipitated with TRPV4 in a dose-dependent manner (Figure 3D,E). These results indicate that TRPV4 does not bind directly to the HH domain of CD98hc or β1 integrin tail, but rather to some other region of CD98hc that appears to be exposed via allosteric conformational changes upon its binding to β1 integrin.

### Molecular modeling of β1-CD98hc-TRPV4 interactions

Molecular dynamics simulation (MDS) was then used to identify the sites within CD98hc where TRPV4 binds. As the crystal structure of CD98hc cytoplasmic tail containing the HH domain is not available, PEP-FOLD3.0 MDS software was used to predict the structure of 46 aa adjacent to the HH domain [CD98hc(P121-S166)] *de novo*. *In silico* rigid docking was performed using the known crystal structure of TRPV4 AR domain and the best fit model of CD98hc(P121-S166) using cluspro2.0. Calculation of the frequency that each residue in human TRPV4 AR domain and CD98hc(P121-S166) forms H-bonds in the models analyzed during computational docking experiments (Supplementary Table S1) predicted that the aa residues most critical for CD98hc-TRPV4 complex formation are R224, E218, R219, K310, R316 and Y236 in the AR domain of TRVP4 and D152, E153, E155, K148, K141, K146 and K140 in CD98hc(P121-S166) (Figure 4A, Supplementary Table S1). A similar approach was used to determine the aa residues involved in the binding of the β1 integrin cytoplasmic tail to the HH domain of CD98hc (Figure 4B, Supplementary Table S2), which corresponded to K794, Y795 and E796 of β1 integrin and E168, W179 and R181 in CD98hc.

**Figure 4.**
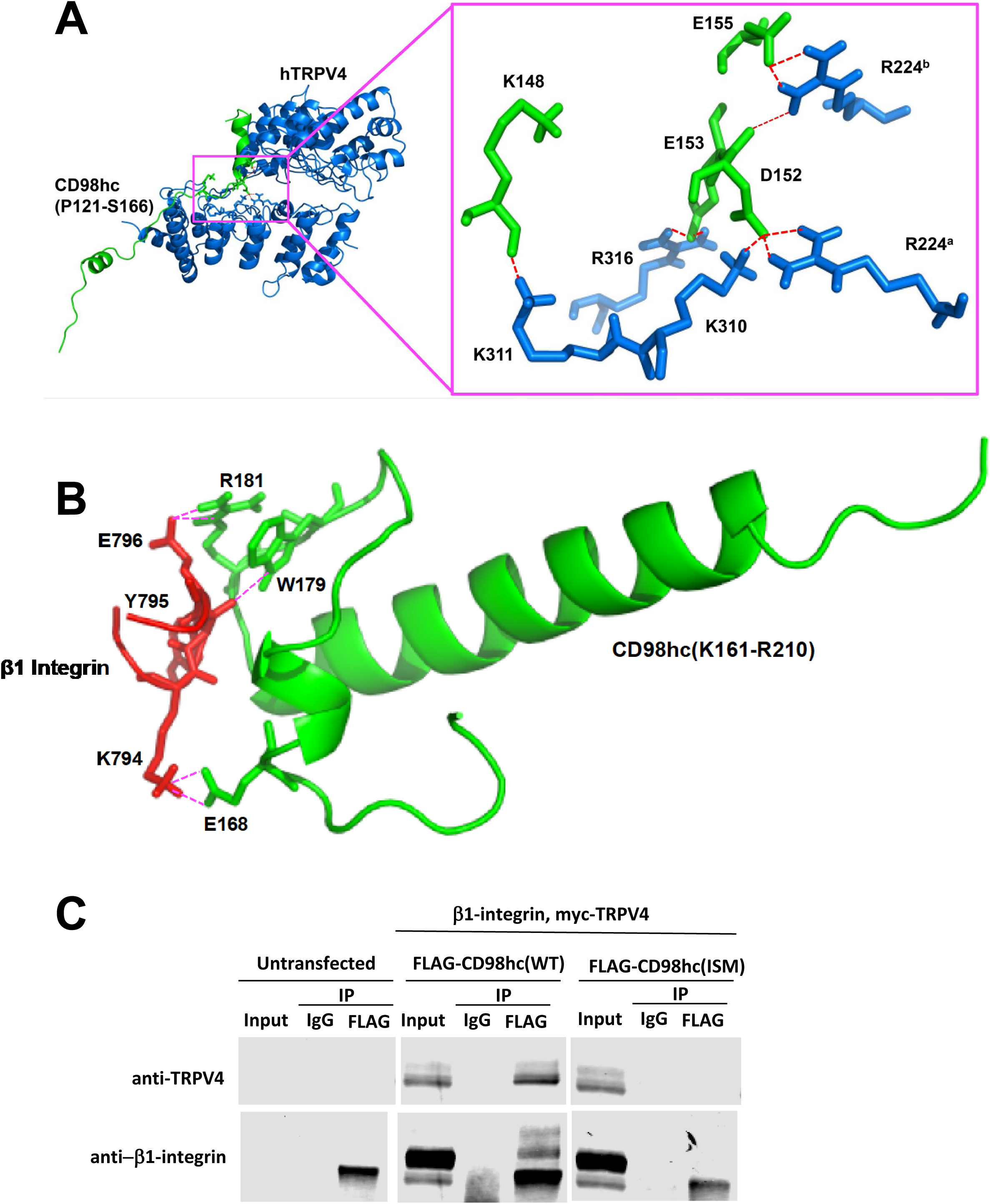
Molecular modeling of β1-CD98hc-TRPV4 interactions. (**A**) Ribbon representation of the predicted complex between human TRPV4 (blue) and the fragment of CD98hc (P121-S166, green). Hydrogen bonds shown as the red broken lines. The crystal structure of human TRPV4 ankyrin was taken from PDB database (PDB ID: 4DX2). Simulation for the cytosolic domain of CD98hc (P121-S166) was performed by PEP-FOLD 3.0. CD98hc (P121-S166) was docked into the 3D structure of hTRPV4 using ClusPro 2.0. Highest scoring model with good topologies is shown. Enlarged stick representation to the right highlights the interacting residues between hTRPV4 and CD98hc (P121-S166). (**B**). Ribbon representation shows molecular modeling of complex between the C-terminus of β1 integrin (N792-K798, red) and the fragment of CD98hc (K161-R210, green). Hydrogen bonds shown as the pink broken lines. Simulations for C-terminus of β1 integrin (N792-K798) and the fragment of CD98hc (K161-R210) were performed using PEP-FOLD 3.0. Integrin (N792-K798) was docked into CD98hc (K161-R210) using ClusPro 2.0. All images were generated using PyMol. (**C**). Immunoblots showing FLAG-tagged CD98hc constructs immunoprecipitated with β1-integrin (bottom) or TRPV4 (top) antibody in lysates from untransfected (left) or HEK293T cells transfected with wild type (middle), mutant CD98hc constructs (right)

CD98hc constructs with specific point mutations were then engineered to experimentally validate these predictions (Supplementary Figure S1B). FLAG-tagged wild type CD98c [CD98hc(WT)] and CD98hc with TRPV4 interacting acidic residues [CD98hc(AA)], TRPV4 interacting basic residues [CD98hc(BA)] or β1 integrin interacting residues [CD98hc(ISM)] mutated were overexpressed in HEK293T cells along with full length β1 integrin and TRPV4. Immunoprecipitation using anti-FLAG antibody and immunoblotting with antibodies against TRPV4 or β1 integrin showed that CD98hc(ISM) failed to pull down TRPV4 or β1 integrin (Figure 4C). The proximity ligation assay also demonstrated that expressing CD98hc containing either mutated TRPV4 interacting acidic or basic residues in HUVE cells significantly decreased (p < 0.0001) the appearance of tight CD98hc-TRPV4 associations that normally appeared evenly distributed across the apical and basal surface of HUVE cells (Figure 5A, B) Thus, these residues appear to mediate close associations between these molecules *in situ*.

**Figure 5.**
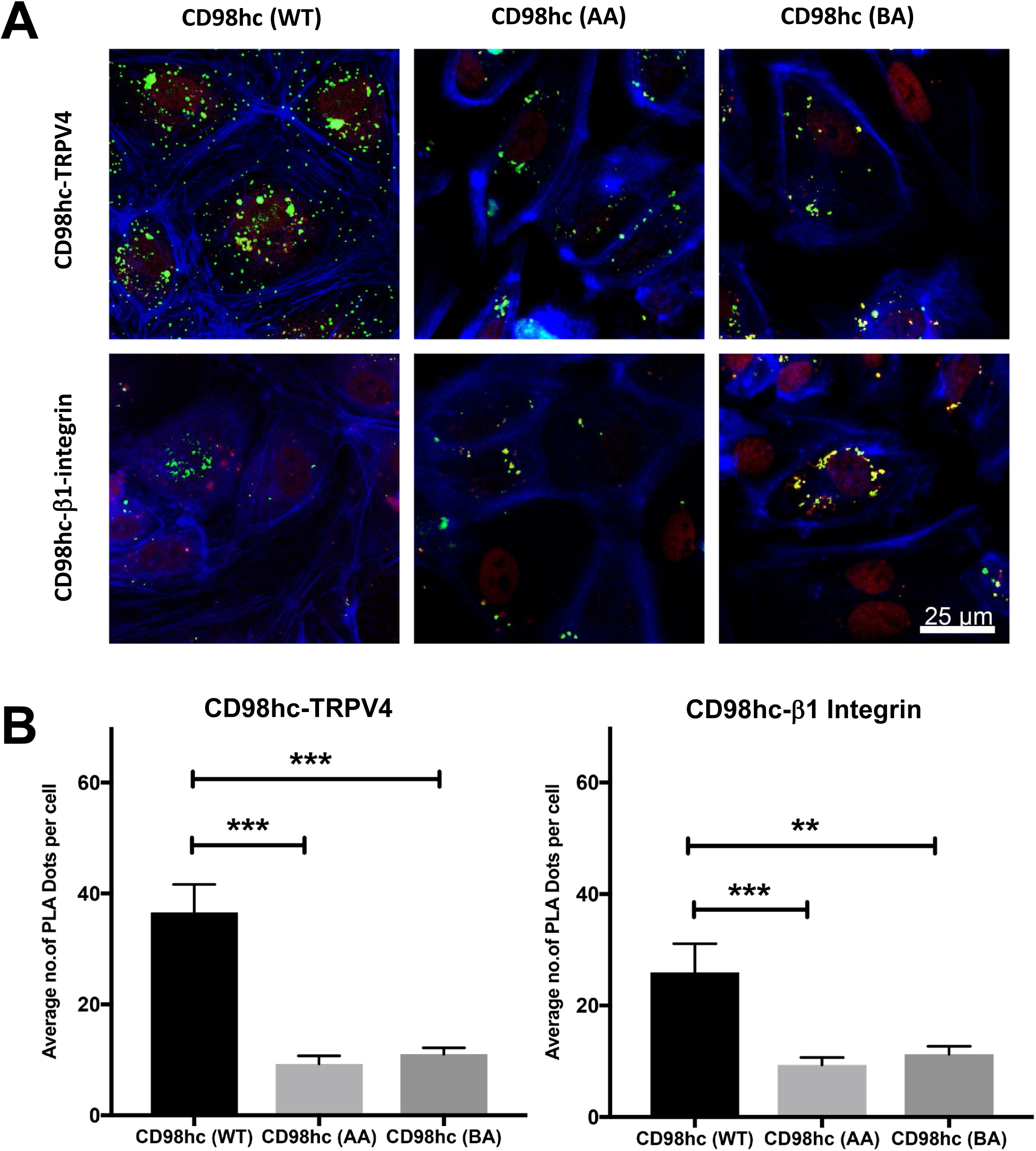
Proximity profiling of β1-CD98hc-TRPV4 interactions. (**A**) Confocal micrographs of basal slices of HUVE cells showing proximity ligation foci (green) between FLAG - CD98hc wild type (WT), mutated acidic residues (AA) or mutated basic residues (BA) and endogenous TRPV4 (CD98hc-TRPV4) or endogenous β1 integrin (CD98hc- β1 integrin) (bar, 20 μm) (**B**)) Histogram showing the average number of PLA dots per cell from experiments described in **A** (** p<0.005, *** p<0.0001).

### CD98hc is required for mechanical activation of TRPV4

To explore the functional relevance of these findings, we knocked down CD98hc using specific siRNA in HUVE cells grown on flexible, ECM-coated substrates (Flexcell dishes) to which mechanical strain (10%) was applied briefly (4 sec). Inhibition of CD98hc expression resulted in reduced activation of calcium influx whereas transfection with a control scrambled siRNA had no effect (Figure 6A,B), as previously described (Matthews et al., 2010). Importantly, inhibiting this mechanically-induced, integrin-dependent, TRPV4 activation mechanism by knocking down CD98hc did not interfere with chemical stimulation of the channel by the known TRPV4 inducer, 4α-phorbol 12, 13-didecanoate (4α-PDD) (Figure 6A,B). Similar specific inhibition of mechanical, but not chemical, signaling through TRPV4 was demonstrated using an RGD peptide that inhibits integrin binding (Supplementary Figure S4). Moreover, both overexpression of the dominant negative CD98hc HH domain (Figure 6C) and knockdown of CD98hc (Figure 6C,D) also altered cell physiology, as indicated by inhibition of the endothelial cells’ normal ability to reorient their shape and internal actin cytoskeleton in a perpendicular direction when exposed to uniaxial cyclic mechanical strain, which was previously shown to be mediated by mechanical activation of TRPV4 through β1 integrin (Thodeti et al., 2009). Hence, CD98hc’s role in mechanically-induced activation of calcium signaling through TRPV4 and regulation of endothelial cell physiology correlates directly with its ability to physically associate with TRPV4 through its cytoplasmic tail, and neighboring sites within this same cytoplasmic region of the CD98hc molecule that also mediates its recruitment into β1 integrin-containing focal adhesions (Supplementary Figure 1B).

**Figure 6.**
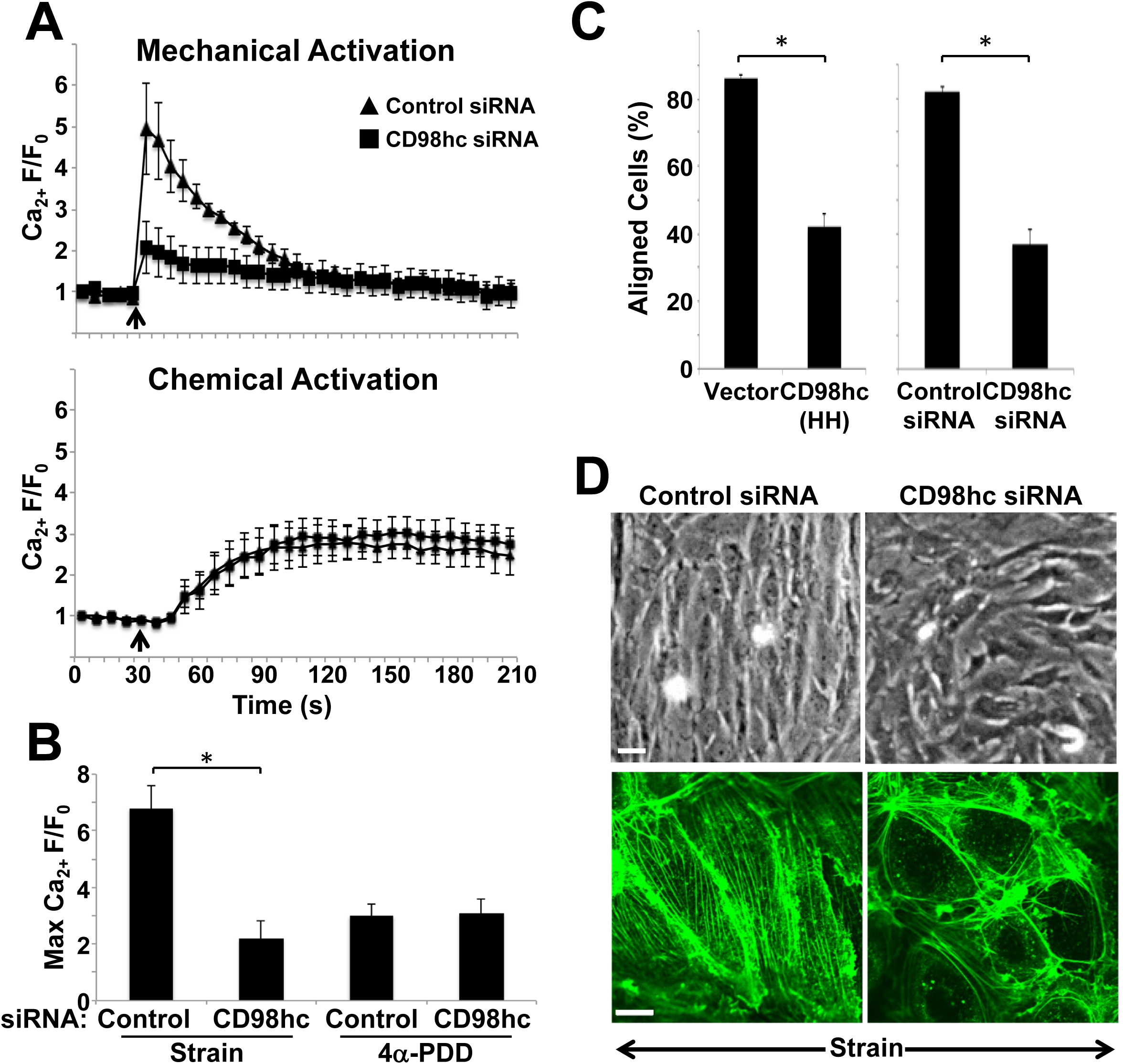
CD98hc is required for mechanical activation of TRPV4. (**A**) Top graph shows relative changes in cytosolic calcium (Ca_2+_ F/F_0_) in Fluo-4-loaded HUVE cells treated with control or CD98hc siRNA in response to static 15% strain for 4 s (arrow indicates time of force application). Lower graph shows the effects of addition of the TRPV4 activator, 4α-PDD (10 mM; arrow indicates time of addition) using the same assay. (**B**) Average maximum relative changes in cytosolic calcium (Max Ca_2+_ F/F_0_) in cells described in **A** (* p<0.05). (**C**) Histogram showing quantification of aligned cellsell (oriented 90 ± 30° relative to direction of stretch) under control conditions or when transfected with CD98hc(HH) or CD98hc siRNA (* p<0.05). (**D**) Phase contrast and fluorescence micrographs of cells stained for F-actin with Alexa 488-phalloidin showing cell reorientation in HUVE cells treated with control or CD98hc siRNA in response to cyclic strain (15%, 1 Hz, 2 h). Arrows indicate the direction of applied strain; bars, 50 μm (top) and 5 μm (bottom).

This form of transmembrane mechanotransduction appears to be direct based on the past demonstration of the rapidity of mechanical activation of TRPV4 by forces applied to integrins (< 4 msec) and that calcium influx localizes to the focal adhesion where direct force is applied (Matthews et al., 2010). This possibility is also supported by our finding that integrin, CD98hc, and TRPV4 bind directly together to form hetero-protein complexes within focal adhesions, but TRPV4 channel activiation is also known to be modulated by its phosphorylation state (Fan et al., 2009). Interestingly, we found that mechanical deformation of the ECM substrate on which HUVE cells are cultured leads to increased phosphorylation of TRPV4, and that this can be inhibited by treatment with CD98hc siRNA (Supplementary Figure S5). Thus, mechanical forces transferred across the integrin-CD98hc-TRPV4 axis may also modulate TRPV4 channel activity through transfer of mechanical forces that induce a conformational change in TRPV4 which makes it susceptible to modification by an associated kinase. Alternatively, TRPV4 may need to be present within focal adhesions for this phosphorylation to take place, or if this kinase always tightly associates with TRPV4, then mechanical force transfer to the kinase via tightly associated TRPV4, CD98hc, and β1 integrin could also result in conformation changes in the kinase that increase its activity and hence, its ability to phosphorylate TRPV4.

## DISCUSSION

Cells contain a tensionally-integrated cytoskeleton that interacts physically with the ECM using specific mechanoreceptors, including β1-integrins(Ingber, 1997; Wang et al., 1993). Mechanical stresses can therefore determine the shape of cell and nucleus (Maniotis et al., 1997) and control numerous cellular functions, including growth, differentiation, migration, and apoptosis (Chen et al., 1997) as well as stem cell lineage commitment (Ingber, 2005; Mammoto and Ingber, 2009; Mammoto et al., 2011; Moore et al., 2002; Sanchez-Esteban et al., 2006). Cellular mechanotransduction involves integrin-mediated force transmission between cells and ECM, which results in force-induced structural changes in proteins that results in mechano-chemical conversion and triggers subsequent mechanical signaling pathways (Geiger et al., 2009; Hoffman et al., 2011). While past studies have demonstrated that mechanical forces transmitted across the cell surface induce deformations in proteins, such as talin and vinculin, within focal adhesion (Choquet et al., 1997), little is known about how the molecular mechanism by mechanical forces applied to ECM activate ion channels on the cell surface, and thereby trigger mechano-electrical conversion processes that are important for control of cell, tissue, and organ physiology. Thus, the major advance of this study is that we have mapped out the path of mechanical signal transfer across cell surface β1 integrins and to TRVP4 ion channels that are activated almost immediately (< 4 msec) after force application to ECM adhesions (Matthews et al., 2010).

Our results show that β1-integrin, CD98hc and TRPV4 colocalize and tightly associate within spontaneously formed mature focal adhesions at the base of the cell, which is consistent with past work showing that they also co-associate in nascent focal adhesions formed at the interface of ligand-coated beads that bind to cell surface β1-integrin receptors (Matthews et al., 2006; Matthews et al., 2010; Wang and Ingber, 1995). However, while intact CD98hc was required for TRPV4 colocalization to focal adhesions, it was not necessary for formation of these adhesions. Moreover, our proximity ligation assay results similarly showed that close β1-integrin-CD98hc interactions are restricted to focal adhesions, whereas CD98hc and TRPV4 associate tightly at sites distributed across the entire cell membrane. These findings suggest that β1-integrin availability is the limiting factor that controls recruitment of CD98hc-TRPV4 complexes into focal adhesions, and formation of this 3-protein hetero-complex.

We also found that the transmembrane and cytoplasmic juxtamembrane domains of CD98hc mediate CD98hc binding to β1 integrin. This is consistent with a past study which similarly showed that the transmembrane domain of CD98hc is sufficient to mediate its binding to β1 integrin, as well as associated induction of PI3K-dependent cell adhesion and migration (Cai et al., 2005). However, we extended these findings by demonstrating that the 50 AA HH domain within CD98hc is responsible for CD98hc’s ability to bind to the β1 integrin tail, and by identifying aa W179 and R181 as the key residues that mediate this interaction using both computational MDS and site-directed mutagenesis. In addition, we discovered that the TRPV4 AR domain is required for CD98hc-TRPV4 binding; however, a dominant negative effect was observed when the CD98hcHH was overexpressed, indicating that TRPV4 binds elsewhere within CD98hc molecule. Importantly, this led to the identification of E153, K148, D152, K141, K146, E155 and K161 as the key aa residues in the cytoplasmic portion of CD98hc that mediate its binding to TRPV4 using in silico docking studies combined with site-direct mutagenesis, co-immunoprecipitations, and proximity labelling assays. These residues were predicted to interact with R224, E218, R219, K310, R316 and Y236 residues in TRPV4, which mapped to a pocket within the second AR domain. This pocket is close to the site where PI(4,5)P_2_ binds and induces rearrangements in the cytosolic tail of TRPV4, thereby facilitating its activation by heat and osmotic stimuli (Garcia-Elias et al., 2013), whereas it is distinct from the site in the AR domain where binding of PI(4,5)P_2_ binding has been shown to reduce channel activity (Takahashi et al., 2014).

Various stimuli promote TRPV4 channel opening and they utilize multiple signaling pathways and second messenger systems (Vriens et al., 2004), and many chemical ligands that modulate TRPV4 activity target its transmembrane domain in areas close to the pore-forming region (White et al., 2016). In contrast, our results suggest that mechanical stresses alter TRPV4 activity by inducing conformational changes in the cytoplasmic AR domains of the molecule, which also represent sites where other physical stimuli, such as heat and osmotic changes, promote TRPV4 activation (Garcia-Elias et al., 2013). The ultra-rapid (< 4 msec) activation of TRPV4 by forces applied to integrins appears to rule out the involvement of soluble second messenger systems that appear to be upstream mediators of TRVP4 activation by heat and osmotic stress (Gao et al., 2003; Garcia-Elias et al., 2013). This is also consistent with our finding that chemical activation of TRPV4 by PDD did not require CD98hc, further emphasizing the mechanospecificity of this β1 integrin-CD98hc-TRPVR signaling axis.

The precise biophysical mechanism by which mechanical force application to β1-integrin and its transfer to TRPV4 via CD98hc results in increased calcium influx through the TRPV4 channel within focal adhesions is not known. Alterations of mechanosensitive ion channel activity commonly result from changes in the 3D conformation of the channel that alters its opening and closing kinetics (Ranade et al., 2015), and this appears to be sensitized (further enhanced) by kinase-induced phosphorylation of TRPV4 (Fan et al., 2009). Here we show that mechanical forces are transferred directly between integrin, CD98hc, and TRPV4 producing conformational changes that may directly influence TRPV4 channel activity, which is consistent with the rapidity of this response (< 4 msec). We also show that knocking down CD98hc also inhibits TRPV4 phosphorylation and thus, either force transfer to TRPV4 or its recruitment to focal adhesions are required for this kinase-mediated enhancement response. One of the many kinases that associate with the cytoplasmic portion of β1 integrin (e.g., FAK, Src, ILK, etc.) in focal adhesions could phosphorylate TRPV4 in response to mechanical strain application, but the kinetics of FAK and Src activation are much slower in response to mechanical force application to integrins than activation of TRPV4, and TRPV4 was shown to be upstream of FAK activation (Thodeti et al., 2009). If ILK or another integrin-associated kinase mediates this activation, then it must be triggered directly by mechanical deformation of integrins or one of its associated molecules given the incredible rapidity of this response. Serine/threonine kinases also could be involved as PKC and PKA have been similarly shown to enhance activation of the TRPV4 ion channel by inducing its phosphorylation, and this depends on assembly of PKC and PKA into a signaling complex with TRPV4 (Fan et al., 2009). This is not involved in direct mechanical activation of TRPV4, but rather only in amplification of this response once triggered mechanically.

The direct mechanism of force transfer between surface receptor (β1 integrin), bound linker protein (CD98hc), and mechanochemical transducer molecule (TRPV4) within the focal adhesion explains why this integrin-specific mechanotransduction mechanism is so fast, occurring within a few milliseconds rather than the seconds or minutes observed for most other integrin-associated mechanochemical conversion mechanisms in non-specialized (non-mechanosensor) cells. As force-dependent activation of TRPV4 depends on mechanical strain and not isometric tension, this arrangement also explains why partially decreasing the pre-stress (isometric tension) in the cytoskeleton can enhance force-dependent distortion of this multi-protein complex (Matthews et al., 2010), and hence, how cytoskeletal tension can ‘tune’ the ultimate mechanochemical conversion response(Mammoto and Ingber, 2009; Wang et al., 2009)

Our finding that CD98hc siRNA and integrin antagonists inhibit TRPV4 activation induced by mechanical strain application through cell-ECM adhesions, but *not* by the chemical activator 4α-PDD that stimulates TRPV4 independently of PKC (Vriens et al., 2004), suggests that these two TRPV4 activation pathways are distinct. This raises the possibility of developing mechanotherapeutics that specifically interfere with mechanical induction of TRPV4 without inhibiting its activation by physiological chemical cues, thereby producing more effective and specific therapeutics for treatment of various diseases, including pulmonary edema.

## METHODS

### CONTACT FOR REAGENT AND RESOURCE SHARING

Please contact Dr. Donald Ingber for further information and requests for resources and reagents.

### EXPERIMENTAL MODEL AND SUBJECT DETAILS

Pooled human umbilical vein endothelial (HUVE) cells were obtained from Lonza, and HEK293T cells were obtained from American Type Culture Collection (ATCC). HUVE cells were cultured in endothelial growth medium supplemented with 5% fetal bovine serum (FBS) and growth factors (EGM-2; Lonza). HEK293T cells were cultured in Dulbecco’s modified Eagle’s medium (DMEM) supplemented with 10% FBS and 100 units/ml penicillin.

### METHOD DETAILS

#### Plasmid construction

CD98hc-HA, CD98hc(ΔHH)-HA, and CD98hc(HH) were constructed using RT-PCR with cDNA from HUVE cells, and subcloned into pCMV-HA vector (Clontech) at the SalI/XhoI sites. TRPV4-myc, TRPV4(ΔPR)–myc and TRPV4(ΔAR1-3)–myc were constructed using PCR with template plasmids from Open Biosystems subcloned into pCMV-myc vector (Clontech) at the SalI/XhoI sites. FLAG tagged CD98hc(WT) was constructed by cloning a synthesized gene block containing the sequence corresponding to 1-210 AA of C98hc and dsredexpress2 connected by a T2A linker into the NotI-EcoRI sites of pcDNA3.1(-)zeo. FLAG tagged CD98hc(ISM), CD98hc(AA) and CD98hc(BA) mutant constructs were constructed by site directed mutagenesis of CD98hc(WT) at specific residues listed in Supplementary Figure. S1B.

#### Protein co-localization experiments

Co-localization experiments were performed on HUVE cells seeded on fibronectin-coated (5 μg/mL, R&D Systems) No. 1 glass bottom dishes (Mattek) and maintained in culture for 24 hrs. Cells were fixed with 4% PFA for 30 min, permeabilized with 0.1% PBS-T for 5min and blocked with blocking buffer (0.03%PBS-T containing 10% Donor donkey serum) for 1h at room temperature. Cells were incubated with above listed combinations of anti-β1 integrin(RRID:AB_2128060 at 1:200), anti-CD98hc(RRID:AB_638284 at 1:50), and anti-TRPV4 (RRID:AB_2040264 at 1:100) overnight at 4C. Cells were washed and stained with appropriate secondary antibodies and imaged using an inverted laser scanning confocal microscope (TCS SP5 X, Leica) using Leica Application Software (LAS, Leica) or using a Zeiss TIRF/LSM 710 confocal microscope equipped with a Hamamatsu ImagEM-1K Back Thinned EMCCD camera controlled by ZEN imaging software (Zeiss).

#### Gene overexpression/knockdown

HUVE cells were transfected using silentFect Lipid Reagent (BioRad) for siRNA studies or Targefect-HUVEC (Targeting systems) for wild type and mutant construct overexpression studies, according to the manufacturer’s instructions. HEK 293T cells were transfected using Lipofectamine 2000 according to manufacturer instructions. Our studies involving siRNA-mediated knockdown were performed in HUVE cells transfected with 50 nM of siRNA against CD98hc or scrambled siRNA duplex (QIAGEN). Wild type and mutant construct overexpression studies were done in HUVE cells seeded on glass bottomed 8-well chamber slides, transfected with 0.5 μg/well of corresponding plasmid DNA and fixed 36h after transfection for immunofluorescent staining and imaging. For gene overexpression studies in HEK293T, cells were seeded in 10cm cell culture dishes, transfected with 10ug/dish of corresponding plasmid DNA and lysed 48h after transfection.

#### Coimmunoprecipitation studies

HUVE cells were extracted in ice-cold Triton buffer (50 mM Tris-hcl, pH 7.4, containing 150 mM NaCl, 1% TritonX100, 5 mM EDTA). Cell extracts were centrifuged at 10,000g for 15 min at 4°C and incubated with magnetic beads (Dynabeads, Invitrogen) conjugated with specific antibodies at 4°C for 2 hours. After washing the beads with Triton buffer, immunoprecipitated protein was detected by immunoblotting. HEK 293T cells were extracted using the above-mentioned ice-cold Triton buffer, sonicated (30% amplitude, 2 pulses of 10sec each), centrifuged, precleared with control IgG beads for 15 min and incubated with anti-FLAG agarose beads (RRID: AB_10063035) at 4°C for 1h. Beads were washed thrice with ice cold Triton buffer and immunoprecipitated protein was detected by immunoblotting.

#### Molecular Dynamics Simulation

The three-dimensional (3D) structures of the C-terminal tail of β1 integrin (N792-K798), the cytoplasmic fragments of CD98hc (P121-S166) and CD98hc (K161-R210) were predicted by PEP-FOLD 3.0 (http://bioserv.rpbs.univ-paris-diderot.fr/services/PEP-FOLD)(Thevenet et al., 2012). In this method, models were clustered by the sOPEP energy value (the coarse-grained energy), and then ranked based on their cluster scores. Among the top 5 ranks, the models with best scores were selected for docking experiments.

#### Molecular Docking Procedure

Docking experiments were performed using ClusPro 2.0 program(Comeau et al., 2004a; Comeau et al., 2004b). The 3D crystal structure of human TRPV4 ankyrin (PDB ID: 4DX2), was obtained from PDB Data Bank. The simulated model of CD98hc (P121-S166) was then docked into the hTRPV4 to generate predicted binding models of CD98hc (P121 –S166) - hTRPV4. The simulated model of the C-terminal tail of β1 integrin (N792-K798) was docked into the cytoplasmic fragment of CD98hc (K161-R210) to generate predicted binding models of β1 integrin (N792-K798) - CD98hc (K161-R210). Models with the highest scores and good topologies were selected for the proposed models of the interaction between TRPV4 and CD98hc, β1 integrin and CD98hc.

#### Proximity Ligation Assay

HUVE cells seeded on IBIDI u-slide were transfected with wild type and mutant CD98hc constructs using Targefect-HUVEC reagent as per manufacturer recommendation. Cells were fixed 36h after transfection with 4% PFA for 30 min at room temperature and proximity ligation assay was performed using mouse or rabbit anti-FLAG, mouse anti-β1 integrin and rabbit anti-TRPV4 antibodies as per manufacturer protocol. Wells were mounted with ProLong™ Glass Antifade and confocal microscopy was performed using Zeiss LSM 880 airyscan or inverted laser scanning confocal microscope (TCS SP5 X, Leica). Images were analyzed using cell profiler program(Carpenter et al., 2006)

#### Wound scratch assay

For the scratch assay, HUVE cells were seeded on fibronectin-coated (5 μg/mL, R&D Systems) No. 1 glass bottom dishes (Mattek) and maintained (2-3 days) in culture and until confluent for 24 hrs. Wounds were created using plastic pipette tips (200 μl, Denville) and images of the scratched monolayers were captured at indicated time points using a Nikon Eclipse TE200 inverted microscope equipped with an RT Monochrome SPOT camera controlled using Spot imaging software (v4.0.9, Diagnostics Instruments Inc.).

#### Calcium imaging

HUVE cells were seeded on stageFlexer-Collagen Type IV-membranes (Flexcellint. Cat# SFM-C-IV-Pack) and 24h later incubated with Fluo-4 dye for 45min at 37°C as per manufacturer recommendations. Cells were washed with live cell imaging solution (Thermofisher Cat#A14291DJ), mounted onto a Flexcell StageFlexer device (Flexcellint. Cat# SF-3000), mechanical strain was applied using Flexcell^®^ FX-4000™ Tension System (Flexcellint. Cat# FX-4000T) and calcium imaging performed using a Nikon eclipse upright microscope fitted with a Hamamatsu EMCCD camera. For mechanical strain induced phosphorylation studies, cells were grown on UniFlex culture plate coated with Collagen Type I (Flexcellint. Cat# UF-4001C) and mechanical strain was applied using Flexcell^®^ FX-4000™ Tension System (Flexcellint. Cat# FX-4000T). Cells were lysed, immunoprecipitated and immunoblotted for phosphorylated proteins using pan anti-Phosphoserine antibody(RRID:AB_11210897)

### QUANTIFICATION AND STATISTICAL ANALYSIS

All values are expressed as the mean ± standard error of the mean (SEM). All experiments were repeated at least 3 times. Statistical comparisons were performed using Student’s t-test for experiments with two conditions or ANOVA for experiments with more than two conditions using GraphPad Prism.

### KEY REAGENTS TABLE

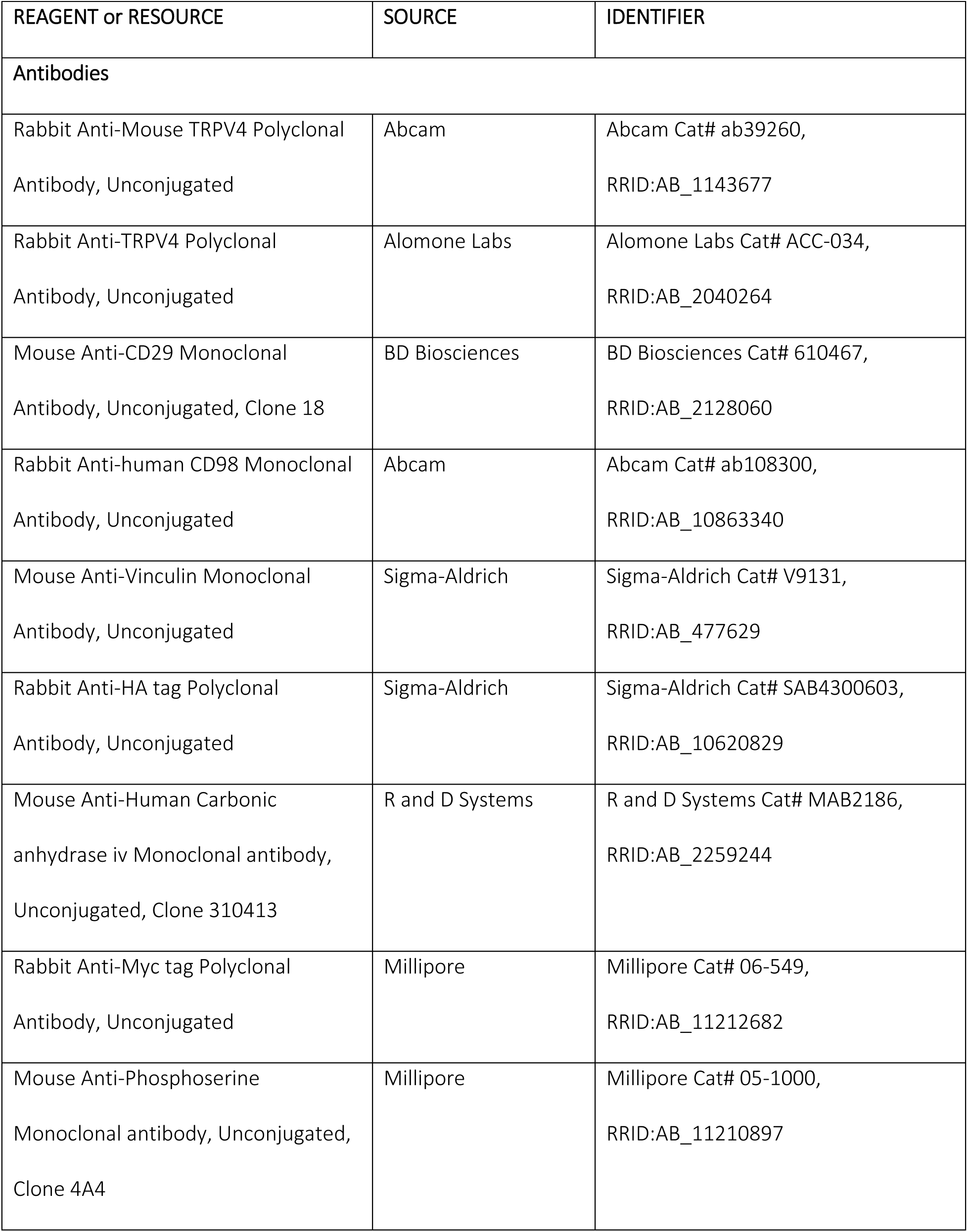
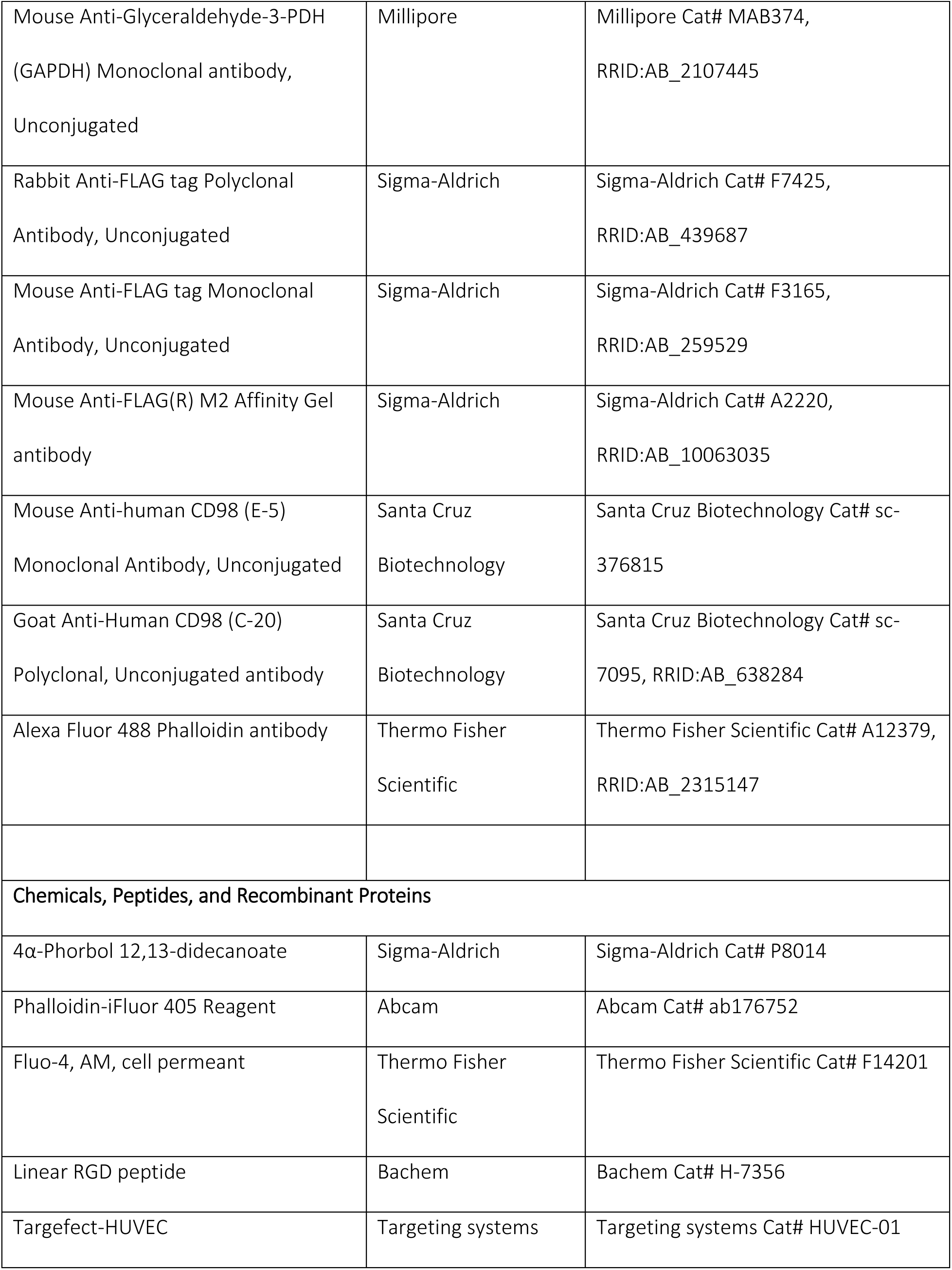
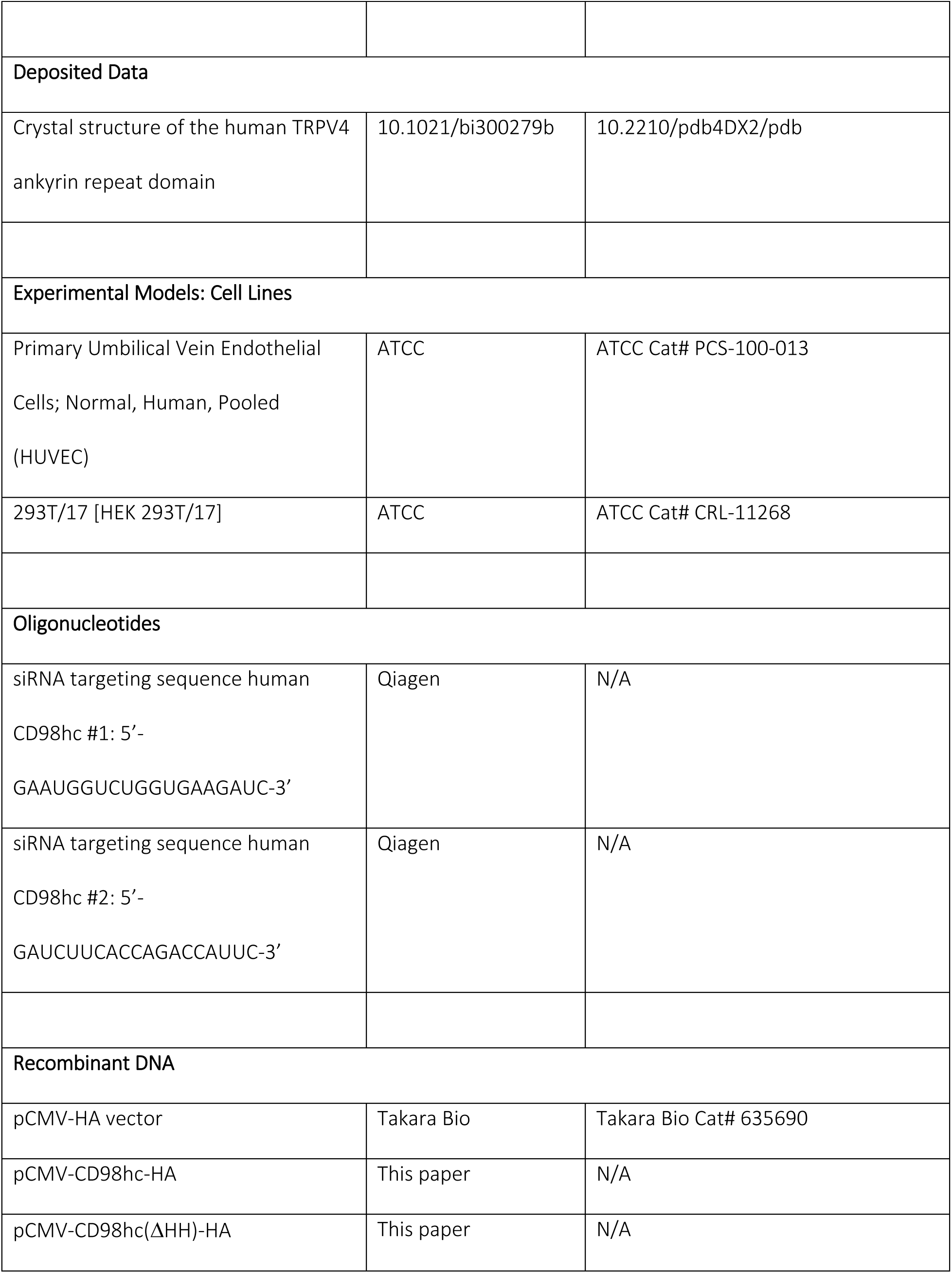
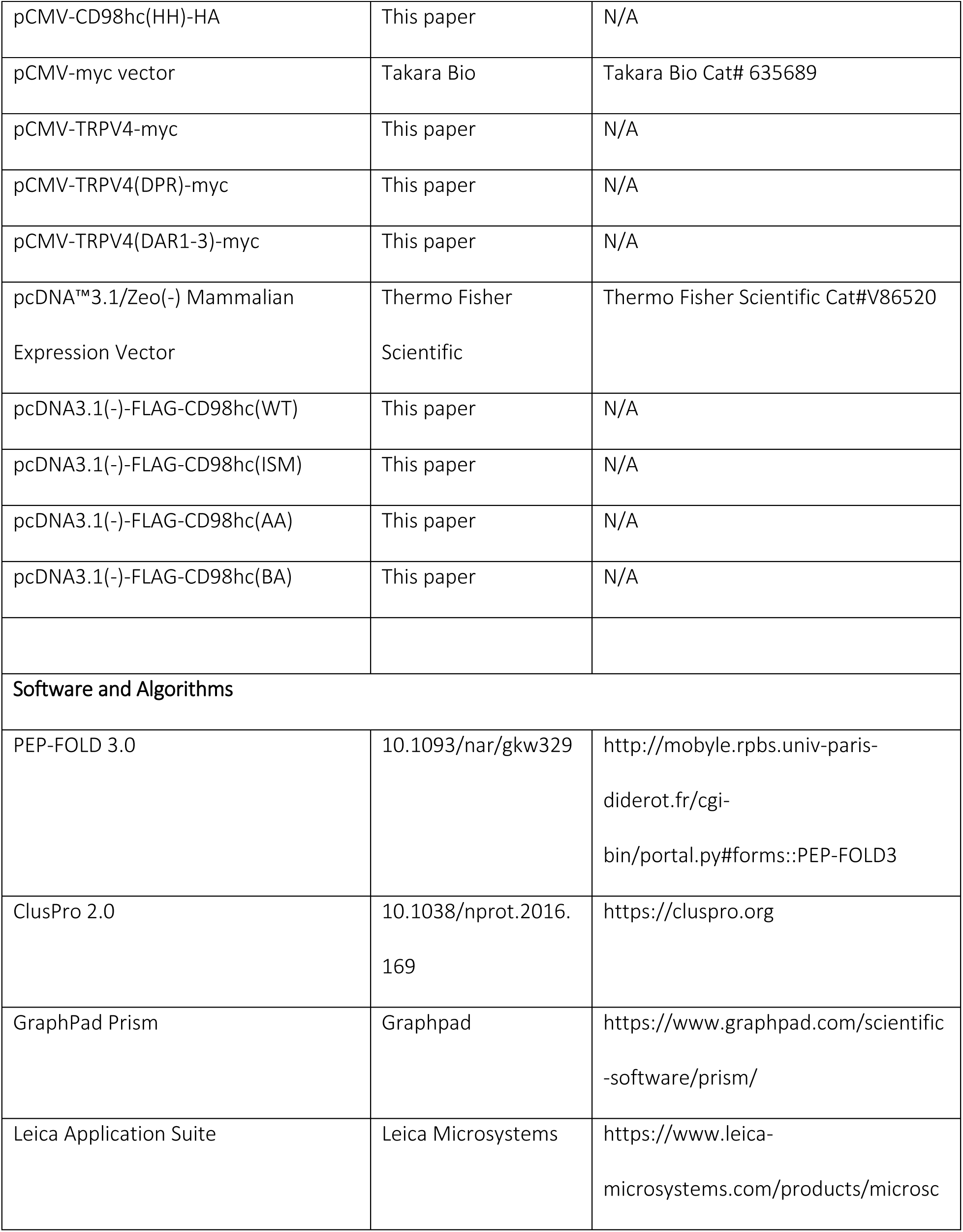
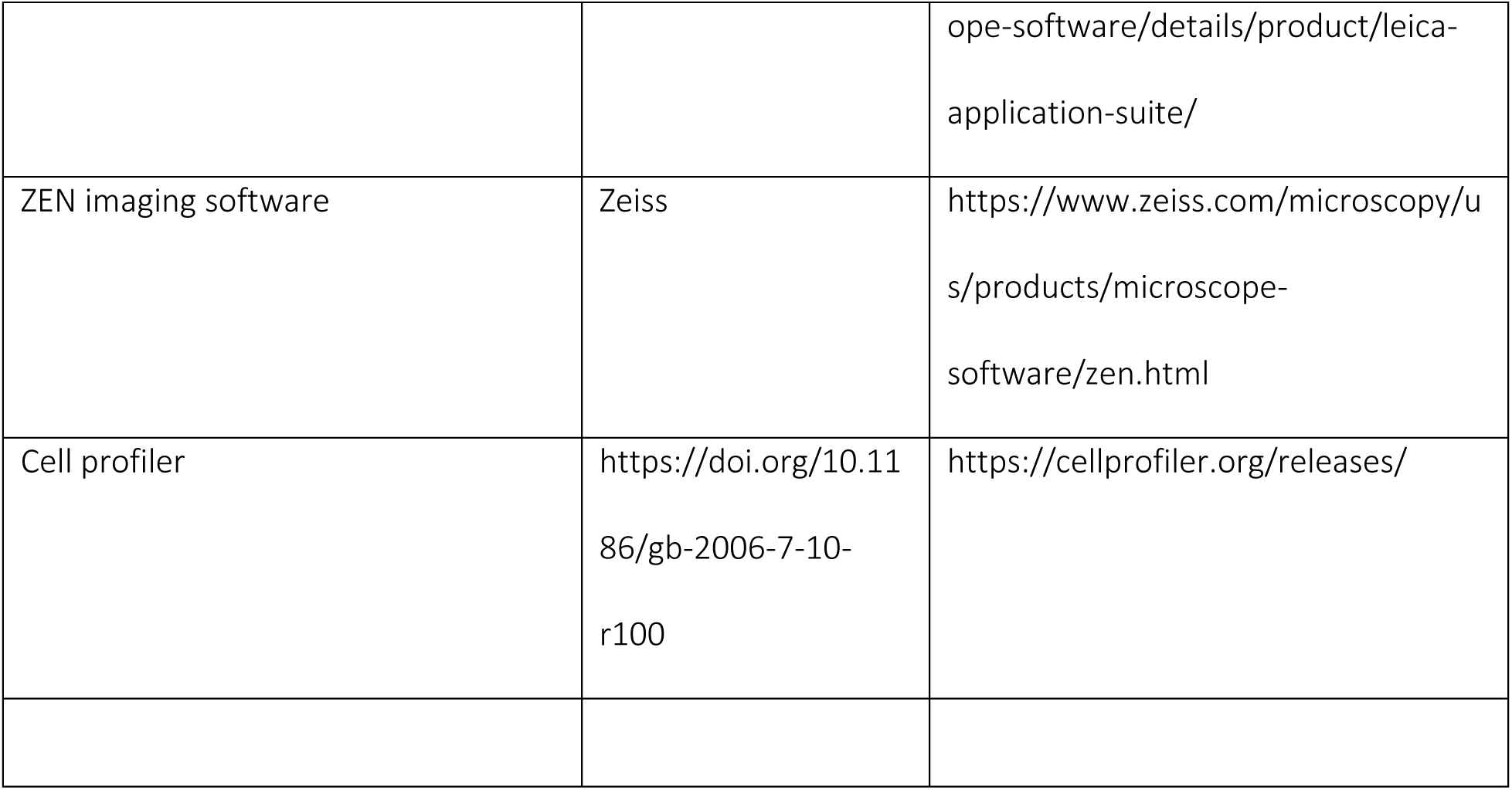

## ACKNOWLEDGEMENTS

This study was supported by a research grant from National Institute for Health (RO1-EB020004) and fellowships from the UEHARA Memorial Foundation and the Japan Society for the Promotion of Science (JSPS). We thank Thomas Ferrante and Sasan Jalili-Firoozinezhad for assistance with TIRF microscopy, Aram Ghalili for assistance with proximity ligation assay and Oren levy for valuable guidance and critic in manuscript preparation.

## AUTHOR CONTRIBUTIONS

Experiments were designed by R.P., M.H.-K., B.D.M., A.M and D.E.I. Experiments were performed by R.P., M.H.-K., H.W., and B.D.M. Data were interpreted by R.P., M.H.-K., B.D.M., A.M. and D.E.I. All authors discussed the results and implications and commented on the manuscript.

---

**Supplementary Figure S1.**
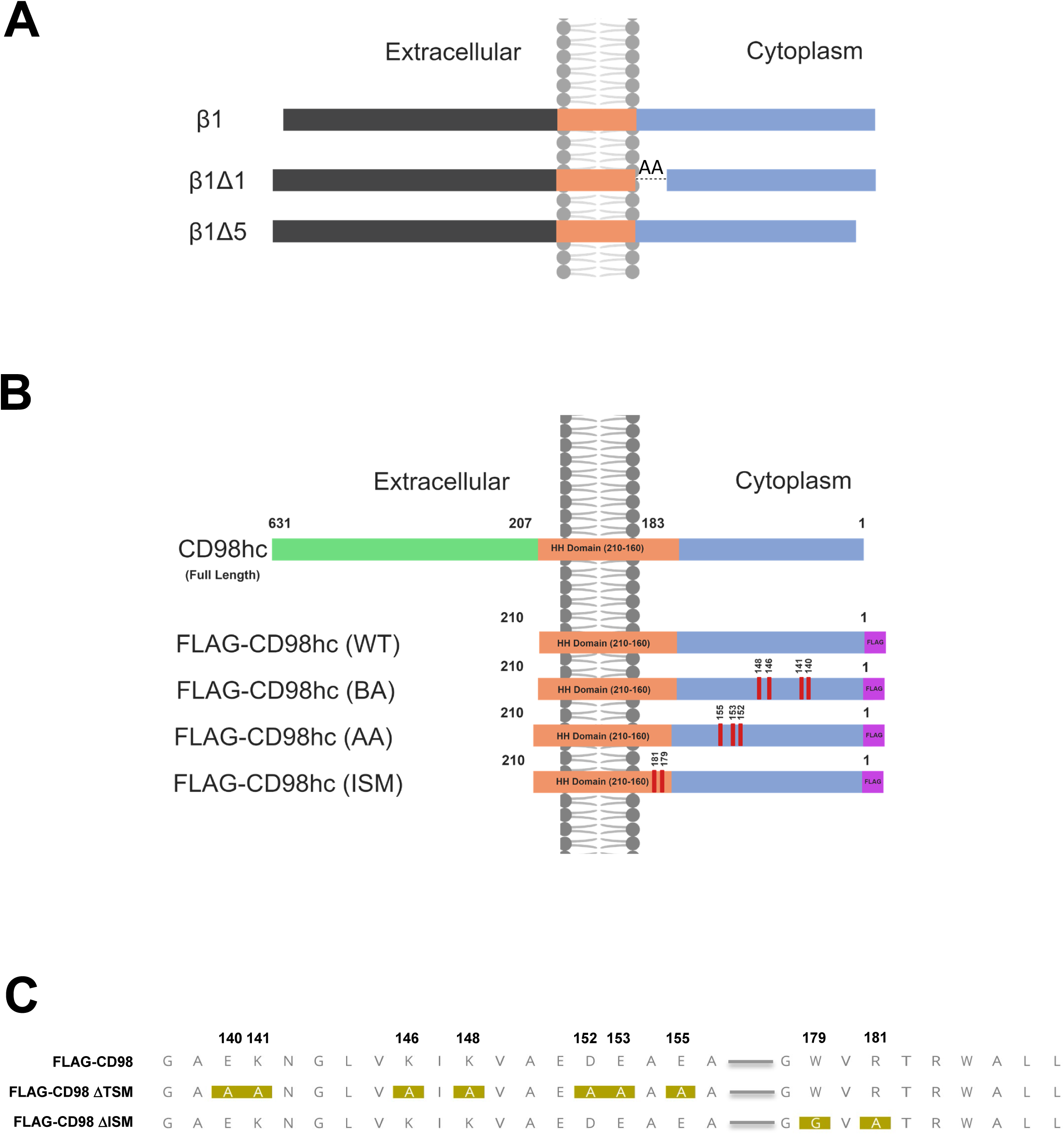
Diagram of mutant β1-integrin and CD98hc constructs. (**A**) Each mutant construct contains the carbonic anhydrase IV (CAIV) enzyme extracellular (EC, black) domain connected to the transmembrane (orange) domain of LDL, and the whole or partial deletions (Δ1 and Δ5) of the β1-integrin intracellular (blue) domain; AA represents replacement of deleted segment with alanine. β1Δ5 represents a construct that lacks last six AA β1-integrin intracellular domain. (**B**) Schematic showing full length CD98hc with its high homology domain (HH domain). Schematic depicting truncated, FLAG tagged CD98hc mutants with wild type (FLAG-CD98hc (WT)), mutated acidic residues (FLAG-CD98hc (AA)), mutated basic residues (FLAG-CD98hc (BA)) and mutated β1-integrin interacting residues (FLAG-CD98hc (ISM)). (**C**) Aminoacid composition of wild type (FLAG CD98), acidic and basic residue mutants (FLAG-CD98ΔTSM) and mutant integrin interacting residues (FLAG-CD98ΔISM).

**Supplementary Figure S2.**
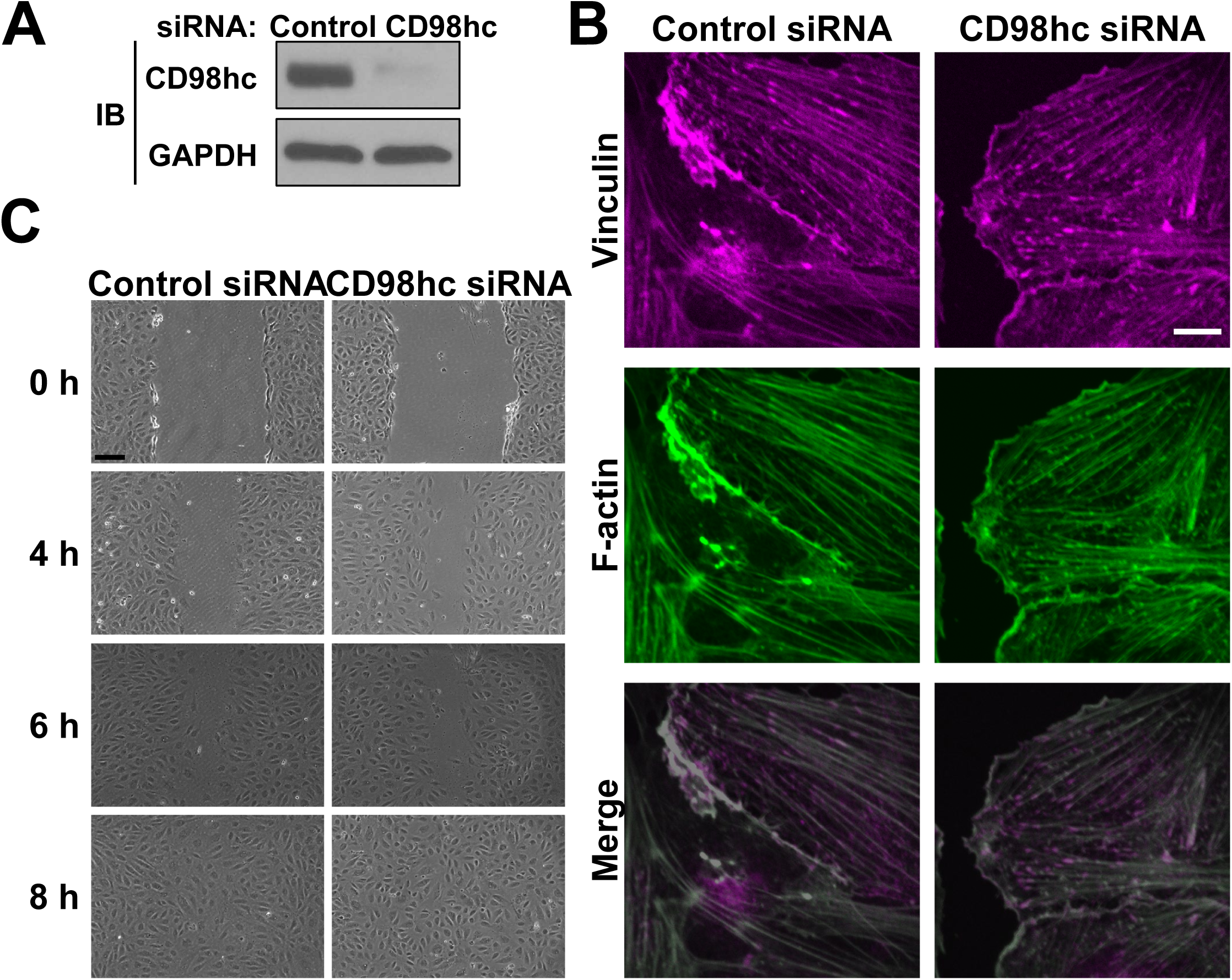
CD98hc is not required for cell migration. (**A**) Immunoblots showing CD98hc and GAPDH protein levels in HUVE cells transfected with control or CD98hc siRNA. (**B**) Fluorescence micrographs of HUVE cells transfected with control or CD98hc siRNA showing formation of focal adhesions containing vinculin (magenta) and F-actin (green). (**C**) Phase contrast photomicrographs showing cell migration following scratch wound of monolayer of HUVE cells transfected with control or CD98hc siRNA (bar, 200 μm).

**Supplementary Figure S3.**
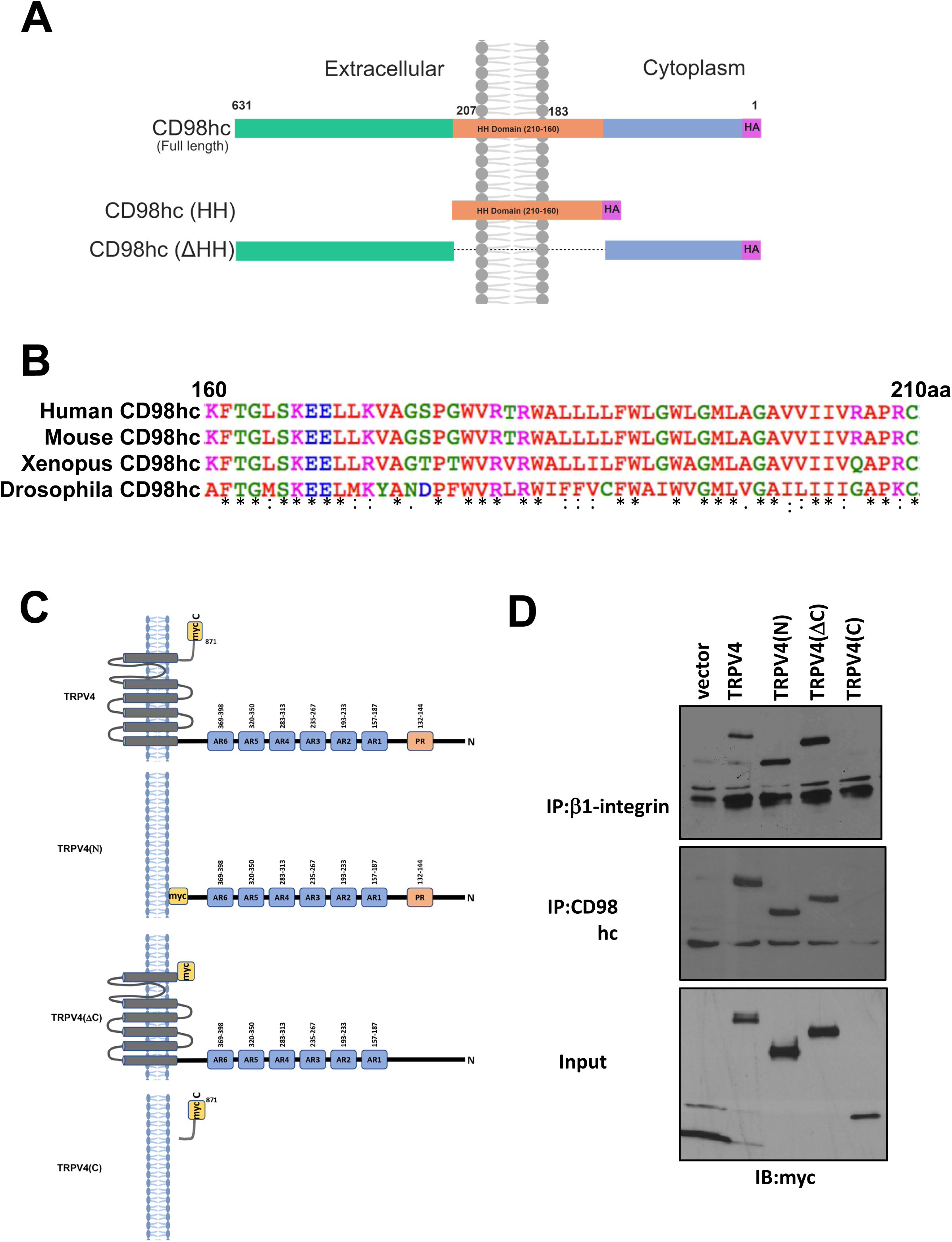
CD98hc HH domain is highly conserved from drosophila to mammals. (**A**) Schematic diagram of CD98hc, CD98hc(HH), and CD98hc(ΔHH) constructs containing high homology (HH, orange) within amino acid (aa) sequences 160-210. (**B**) Amino acid sequences 160-210 in human, mouse, *Xenopus*, and *Drosophila* CD98hc (“*” indicates identity, “:” indicates high similarity, and “.” indicates low similarity in amino acids between the different species).

**Supplementary Figure S4.**
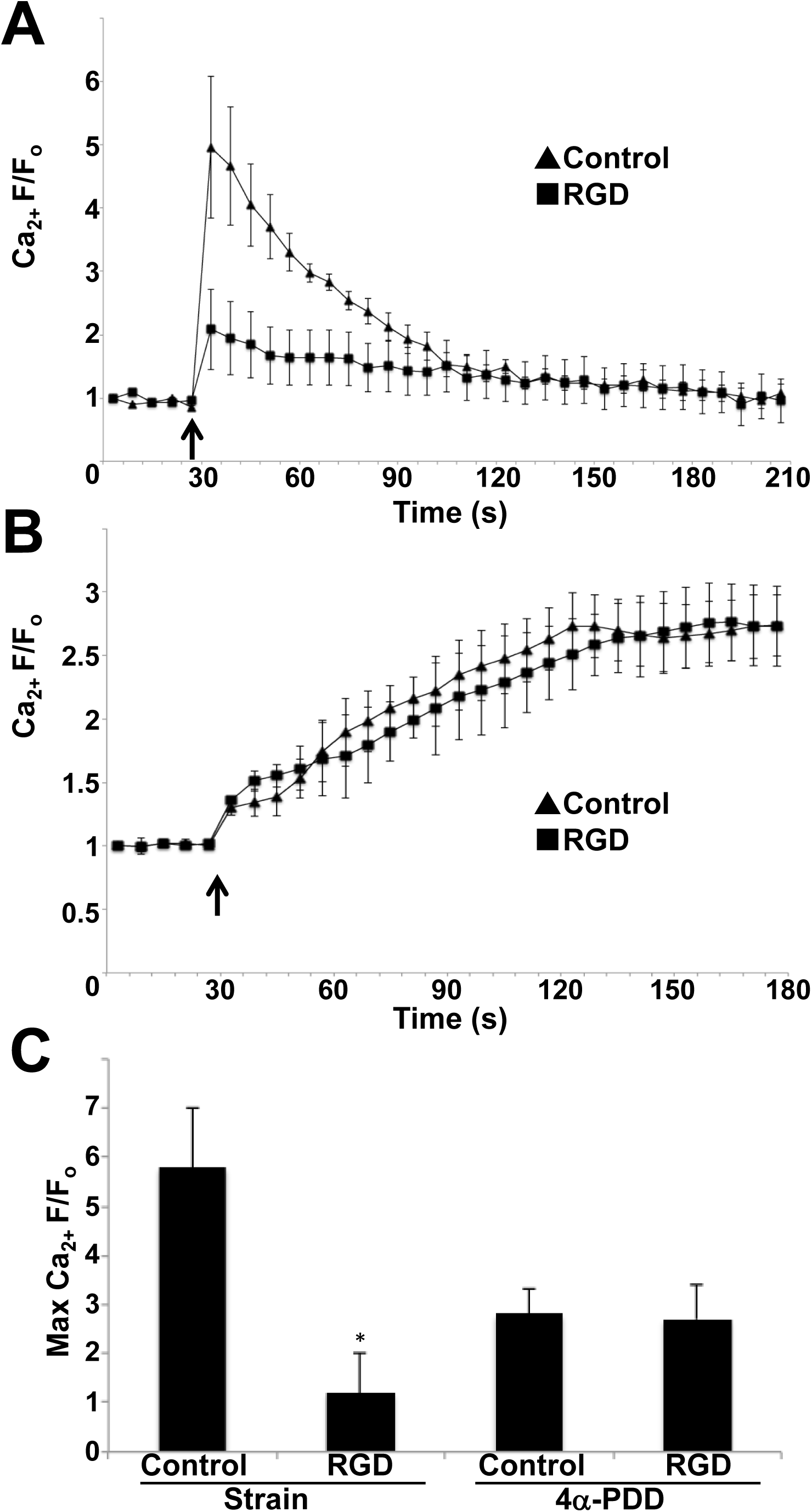
Mechanisms of mechanical and chemical activation of TRPV4 are distinct. Relative changes in cytosolic calcium (Ca_2+_ F/F_0_) in Fluo-4-loaded HUVE cells in response to 15% static strain for 4 s (**A**) or the TRPV4 chemical activator, 10 μM 4α-PDD (**B**) in the presence or absence of RGD peptide (10 μM). (**C**) Average maximum relative changes in cytosolic calcium (Max Ca_2+_ F/F_0_) in cells described in panels **A** and **B** (* p<0.05 vs strain control).

**Supplementary Figure S5.**
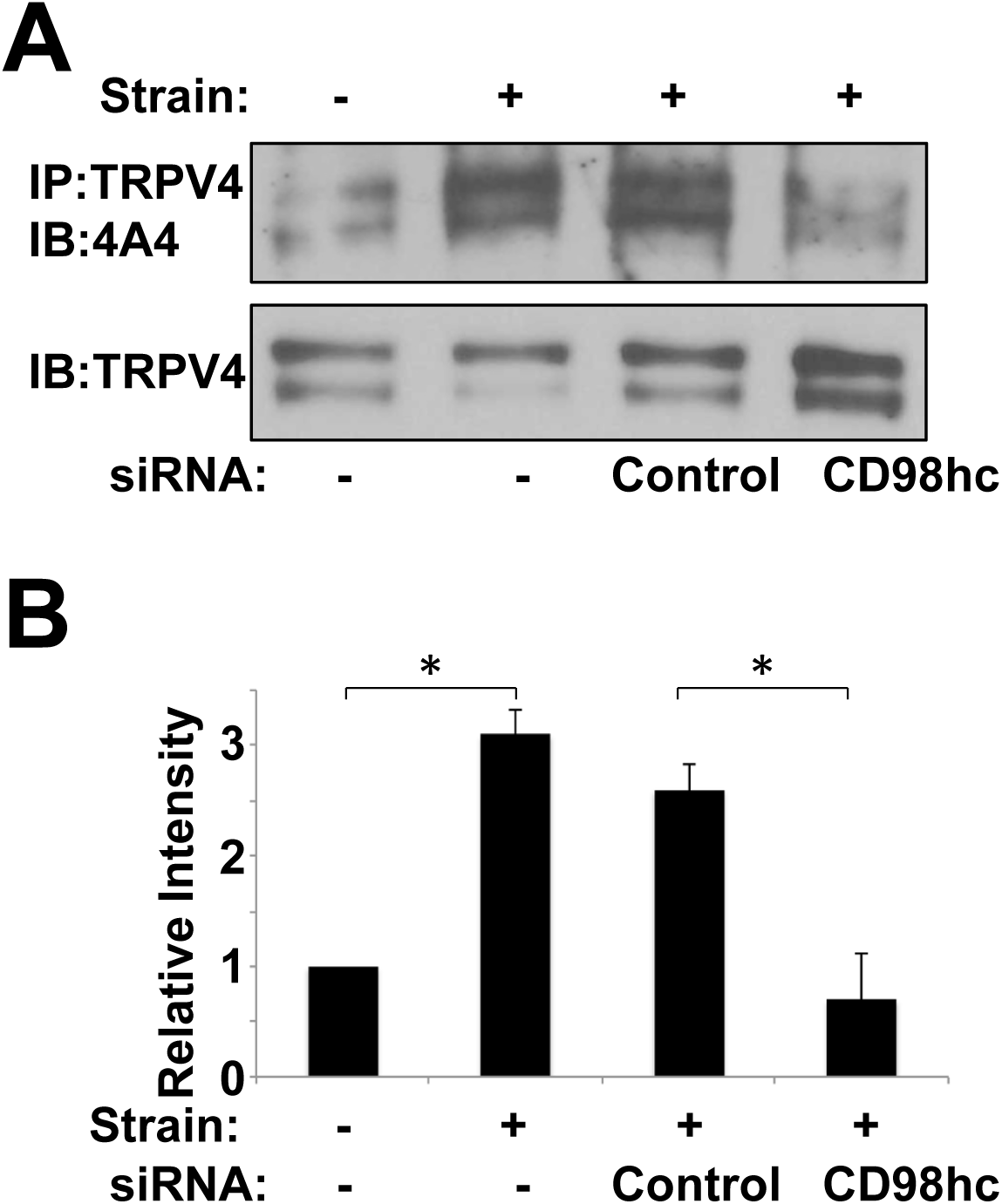
CD98hc knockdown prevents TRPV4 phosphorylation induced by mechanical strain. (**A**) Immunoblots showing phosphorylated TRPV4 from lysates of control HUVE cells or HUVE cells treated with control or CD98hc siRNA and exposed to cyclic substrate strain (15%, 1 Hz, 2 h) detected using the 4A4 anti-phosphoserine antibody. (**B**) Histogram showing the densitometric quantification of immunoblots in **A** (*p<0.05).

**Supplementary Table S1.**
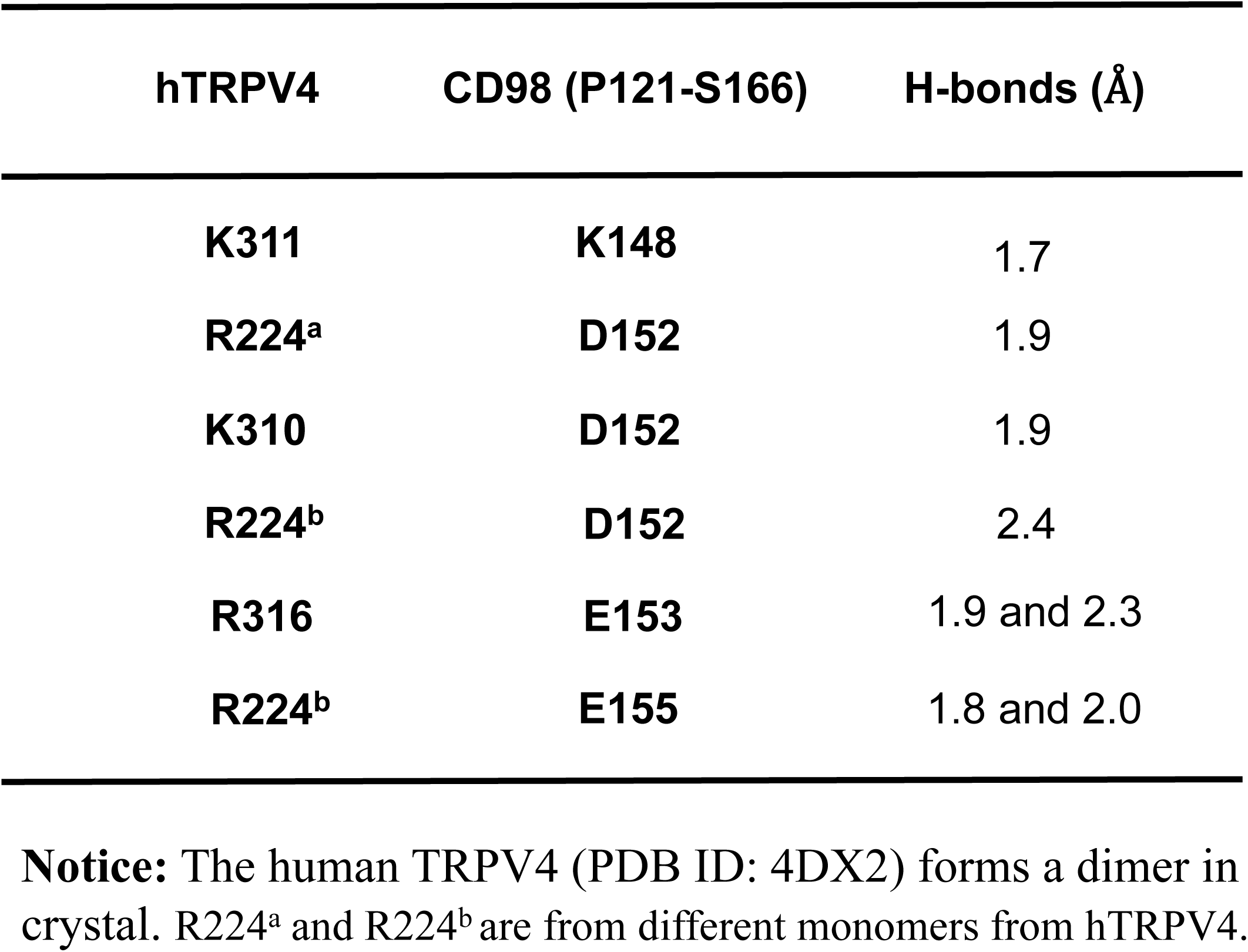
Interaction residues between hTRPV4 and CD98 (P121-S166). Predicted interacting residues in hTRPV4 and CD98 (P121-S166), and corresponding hydrogen (H) – bonds between the residues. **Notice:** The human TRPV4 (PDB ID: 4DX2) forms a dimer in crystal. R224^a^ and R224^b^ are from different monomers from hTRPV4.

**Supplementary Table S2.** Interaction residues between the cytoplasmic tail of β1 integrin (N792-K798) to the HH domain of CD98hc (K161-R210).

| **β1 integrin (N792-K798)** | **CD98hc (K161-R210)** | **H-bonds (Å)** |
| --- | --- | --- |
| **K794** | **E168** | 2.5 and 2.7 |
| **Y795** | **W179** | 2.8 |
| **E796** | **R181** | 2.8 and 3.2 |

